# Cell-type-selective synaptogenesis during the development of excitatory connectivity in the mammalian neocortex

**DOI:** 10.1101/2025.08.08.669371

**Authors:** Alan Y. Gutman-Wei, Sriram Sudarsanam, Alec G. Cabalinan, Naseer Shahid, Anny Shi, Luis E. Guzman Clavel, Sophia M. Spindler-Krage, Amit Agarwal, Alex L. Kolodkin, Solange P. Brown

## Abstract

The function of mammalian neocortex relies on the timing of axon extension and establishment of cell-type-biased patterns of excitatory synaptic connections. A subtype of excitatory neurons, layer 6 corticothalamic neurons (L6CThNs), ultimately exhibit a marked preference for synapsing onto parvalbumin-positive (PV) inhibitory interneurons over more common excitatory cells in layers 6 and 4 (L6, L4). We show that the intracortical axons of L6CThNs develop in phases, elongating within L6, then pausing before extending translaminar branches into L4. Decreasing L6CThN excitability selectively enhanced axon growth in L6 but not later elaboration in L4. For both layers, we tested whether preferential synaptogenesis onto rarer PV interneurons, or promiscuous synapse formation followed by selective pruning, generated adult connectivity. We found that L6CThNs formed functional AMPA-receptor-containing synapses preferentially onto PV interneurons. Silent L6CThN synapses were not detected. Our findings show that cell-type-biased synaptogenesis underlies the formation of functional cell-type-specific excitatory connections in the neocortex.

## INTRODUCTION

Cell-type-biased patterns of synaptic connectivity among neuronal subtypes in the mammalian neocortex underlie its function and contributions to behavior. Recent studies of cortical inhibitory neuron subtypes describe mechanisms that influence their choice of postsynaptic targets during development^1–6^. Different classes of excitatory cortical projection neurons also connect preferentially with specific intracortical postsynaptic partners both within and across cortical layers^1,7–13^, but how these patterns of synaptic connectivity are established during development is not well understood.

Layer 6 corticothalamic neurons (L6CThNs), one of the first types of excitatory pyramidal neuron generated in primary somatosensory neocortex (S1) of mice^14^, provide a striking example of preferential synaptic connectivity. L6CThNs in upper L6 of S1, identified by their long-range axons projecting to the ventral posterior medial (VPM) nucleus of the thalamus, also have intracortical axons that elaborate in L6 and project upward to L4^15–19^. In both L4 and L6, the intracortical axons of L6CThNs form frequent synaptic connections onto parvalbumin-positive (PV) inhibitory interneurons^9,16,20–27^. In contrast, L6CThNs rarely synapse onto excitatory neurons^9,20–24,28–31^, even though excitatory neurons far outnumber PV interneurons in both cortical layers^32,33^. The synaptic preference of L6CThNs for PV interneurons relative to neighboring excitatory neurons is thought to be central to their roles in regulating sensory cortical responses^25–27,34–38^. How the intracortical processes of L6CThNs develop during the first two postnatal weeks, a period of rapid intracortical synaptogenesis^39–42^, and whether they preferentially synapse onto PV neurons during early synapse formation, or instead prune inappropriate synapses onto excitatory neurons to generate the adult patterns of synaptic connectivity, is unknown.

Prior studies show that, in some developmental contexts, long-range axonal projections of pyramidal neurons grow beyond their target areas before being pruned^43–45^. In contrast, many^46–51^, but not all^52^, studies report no net pruning of pyramidal neuron intracortical axons during the first postnatal weeks. In addition, manipulating the intrinsic excitability of pyramidal neurons influences the development of some long-range axonal projections. For example, suppressing the activity of corticocallosal pyramidal neurons via overexpression of the inwardly rectifying potassium channel Kir2.1 limits the development of their axons in the contralateral hemisphere^53–58^. How intrinsic neuronal excitability influences the development of intracortical pyramidal neuron axons is less clear. The timing of the development of the dendritic arbors of L6CThNs relative to their axonal arbors across cortical layers and the influence of neural activity on their development is also not fully understood. Reducing the intrinsic excitability of L2/3 pyramidal neurons via Kir2.1 overexpression suppressed the development of their dendritic arbors^59^, but whether this effect generalizes across pyramidal cell types is not known.

Are patterns of excitatory synaptic connectivity in the cortex established during initial synaptogenesis or through pruning of inappropriate synapses? For inhibitory cortical interneuron classes, which form synapses onto specific cell types and subcellular compartments, both specific synaptogenesis and promiscuous synaptogenesis followed by pruning have been observed during postnatal development^1,3,40,60^. For excitatory L6CThNs, preferential synaptogenesis onto PV interneurons during axon development may generate the adult pattern of connectivity; alternatively, this pattern may emerge only after the preferential elimination of inappropriate synapses onto excitatory neurons. Furthermore, excitatory neurons form silent synapses, glutamatergic synapses with NMDA but no AMPA receptors^61,62^. These silent synapses, which have been detected in neonatal L6^63,64^, may contribute to neural circuit refinement during cortical development by undergoing selective AMPA receptor insertion to generate fully functional synapses^61,62^. Thus, another possibility is that L6CThNs form silent synapses promiscuously, with those onto PV interneurons preferentially undergoing AMPA receptor insertion.

Here, we determine the developmental time course of L6CThN intracortical axon elaboration across cortical layers during the first two postnatal weeks, and the effects of suppressing neuronal activity on their development. Next, we determine whether L6CThNs synapse preferentially onto PV interneurons to generate their adult patterns of synaptic connectivity. Our results represent important steps in understanding the formation of cell-type-biased excitatory connectivity in the cortex.

## RESULTS

### Intracortical axons of layer 6 corticothalamic neurons develop in two postnatal phases

To form mature cortical circuits, the intracortical axons of pyramidal neurons must elaborate across cortical layers to come into close contact with their postsynaptic targets. In adult sensory cortex of rodents and other species, L6CThNs elaborate axon collaterals that branch off the primary axon, project upward towards the pia^16,17,19,65–67^, and form synapses both locally within L6 and with postsynaptic partners in L4^9,16,20–31^. Thus, we first determined the timing of L6CThN intracortical axon development during the first two postnatal weeks, a period of considerable intracortical axon development and synaptogenesis^39–42^. We sparsely labeled individual L6CThNs with red fluorescent protein (RFP) using *in utero* electroporation of Cre-dependent constructs in *Ntsr1-Cre* mice, a mouse line which expresses Cre recombinase in L6CThNs in somatosensory cortex^20,68^. We then assessed the processes of electroporated L6CThNs in S1 imaged in intact, cleared, brains using light sheet microscopy followed by full reconstruction of individual intracortical axonal and dendritic arbors from mice at four ages: postnatal day (P)4, P7, P10 and P14 (Figure 1A, 1B, Figure S1). Because there are different types of L6CThNs^16–19,66,67,69,70^, we enriched our analyses for one well-defined subtype: VPM-only L6CThNs. This subset of L6CThNs, identified by long-range axons that project solely to VPM, has cell bodies located in the upper half of layer 6 in rodents^16,18,19,70,71^. We found that both the total intracortical axonal length and the total number of intracortical axon branches of L6CThNs increased in two distinct phases, first increasing significantly between P4 and P7, and then again between P10 and P14 (Figure 1C,1D). Axonal growth slowed significantly between P7 and P10 relative to both the early and later time periods. We did not observe, however, net axon pruning during the first two postnatal weeks, consistent with prior studies of other pyramidal cell types in rodents^46,48–51^. Thus, during the first two postnatal weeks, the overall axonal arbors of L6CThNs undergo two major phases of intracortical axon outgrowth, separated by a pause.

**Figure 1.**
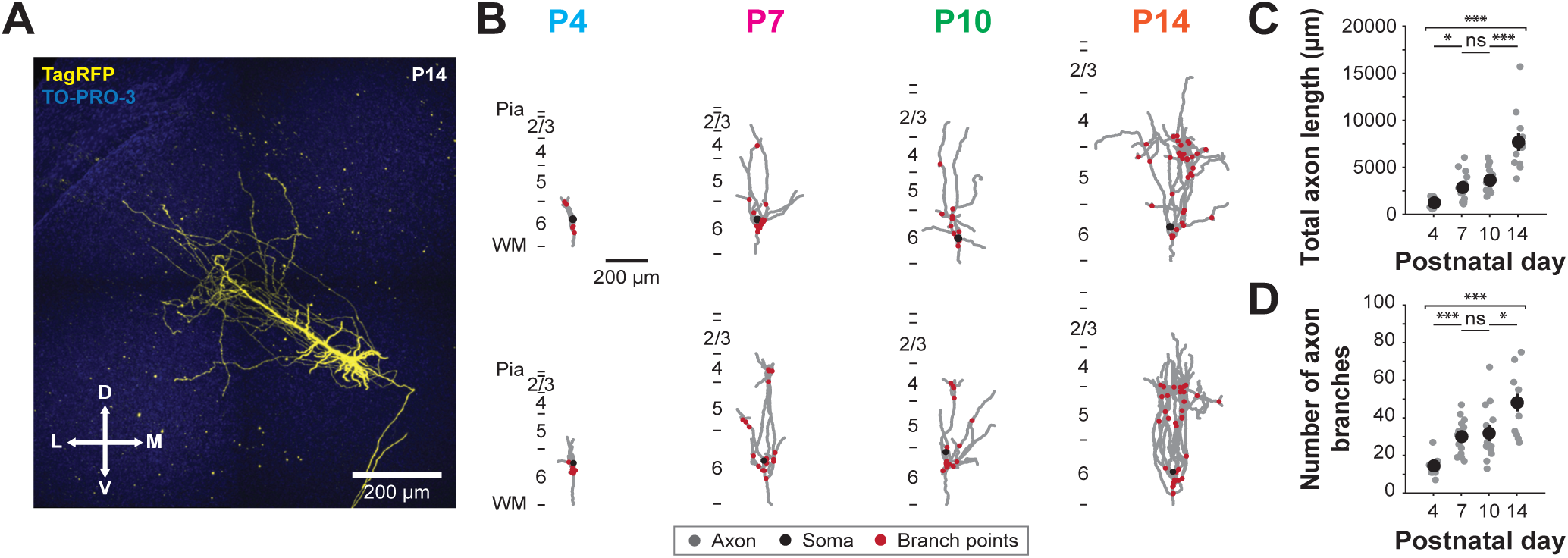
The intracortical axons of layer 6 corticothalamic neurons (L6CThNs) show two major postnatal phases of outgrowth and branching. (A) Example L6CThN labeled with tRFP (yellow) in a cleared brain from a postnatal day 14 (P14) mouse shown as a maximum projection along the anterior-posterior axis. Nuclei were labeled with TO-PRO-3 (blue). (B) Example axonal reconstructions (gray) of L6CThNs from P4, P7, P10, and P14 mice. Red circles represent branch points; black circles represent soma locations. Two example reconstructions are shown per age. (C) Summary data showing the total length of intracortical axon for L6CThNs at P4, P7, P10, and P14. Black circles represent the mean; grey circles represent individual cells (Kruskal-Wallis test, p < 10^-6^; P4: n = 10 cells, N = 3 animals; P7: n = 14 cells, N = 5 animals; P10: n = 12 cells, N = 5 animals; and P14: n = 12 cells, N = 8 animals). (D) Summary data showing the total number of intracortical axonal branches for the same L6CThNs as in C. (Kruskal-Wallis test, p < 10^-4^). C, D: Pairwise Wilcoxon rank-sum tests with Bonferroni correction for multiple comparisons were used to compare between consecutive age groups (*, p < 0.05; **, p < 0.01; ***, p < 0.001). Data in C, D shown as mean ± SEM. Scale bars: A, B: 200 µm.

### The timing of axon development of layer 6 corticothalamic neurons differs across cortical layers

The two major periods of postnatal intracortical axon growth may reflect differences in the timing of L6CThN axon elongation and elaboration across the cortical layers. To compare the layer-specific distribution of L6CThN axons at each postnatal age sampled, we annotated cortical layers as surfaces in the three-dimensional volumes of imaged neurons, segmented reconstructed intracortical axons using these layer boundaries, and quantified L6CThN axon lengths and branch points occurring within L6, L5, and L4 (Figure 2A). In L6, L6CThN axon growth occurred primarily between P4 and P7, with significant increases in axon length and in the number of axonal branch points during this period, but not between P7-P10 or P10-P14 (Figure 2B, 2C). For L4, in contrast, we found that axon development shifted to later ages, with both total axon length and the number of axonal branch points increasing significantly between P10 and P14, but not from P4 to P7 (Figure 2F, 2G). Axon growth in L5 shared features of both L4 and L6. Axon length and branch point number increased significantly both between P4-P7, like L6, and P10-P14, like L4, while not significantly changing during the P7-P10 period (Figure 2D, 2E). Taken together, these results show that L6CThN axon growth occurs progressively across cortical layers, with significant axon elaboration occurring between P4 and P7 in infragranular layers in contrast to L4, where axon length and branchpoint number increase primarily between P10 and P14.

**Figure 2.**
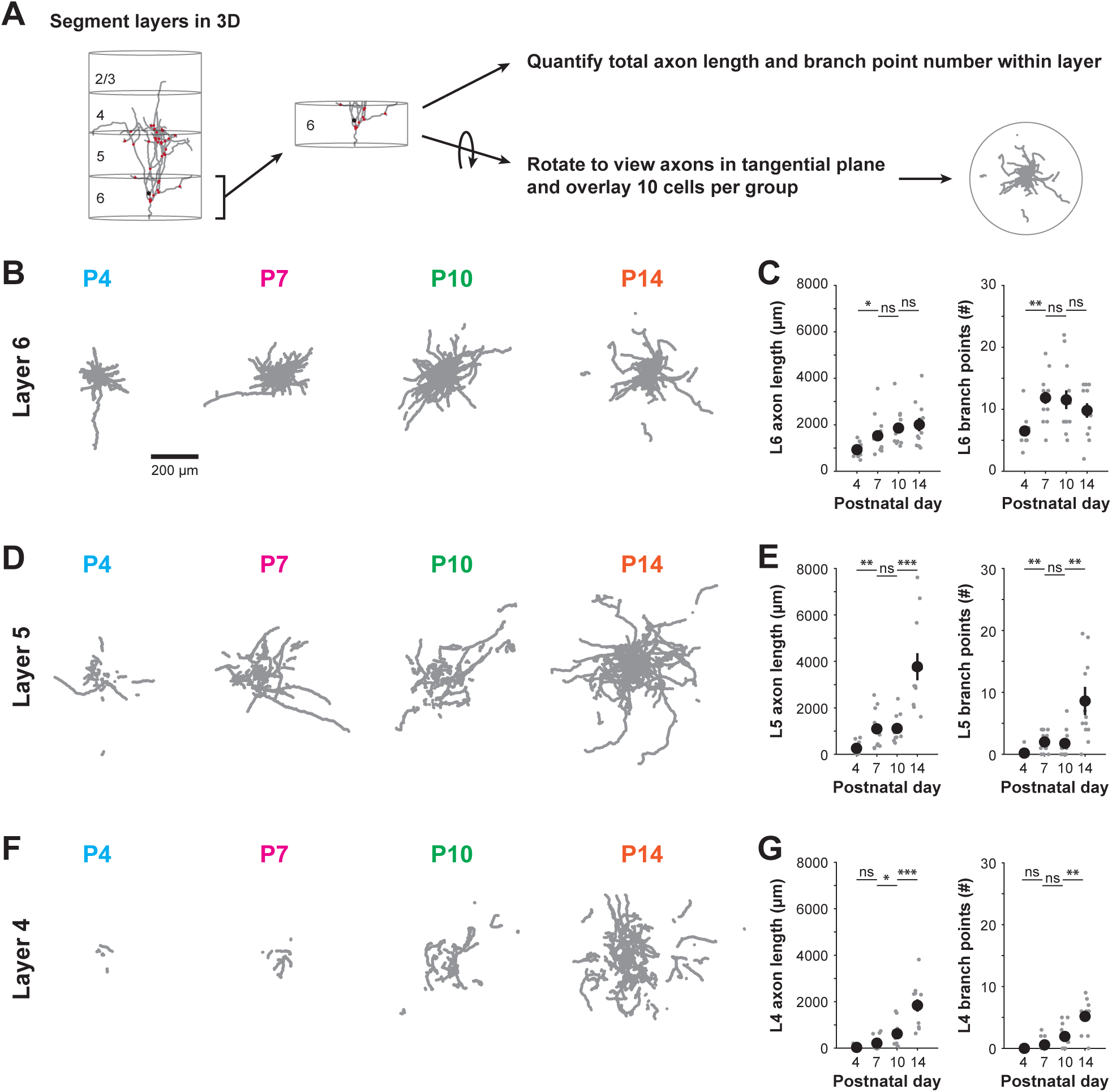
The timing of axon development of layer 6 corticothalamic neurons (L6CThNs) differs across cortical layers. **(A)** Schematic showing the analysis pipeline. For each reconstructed L6CThN, the volume spanned by its axons was segmented into layers and the axonal properties calculated for each layer. For the visualizations in B, D and F, the 10 cells with axon lengths within each layer closest to the mean of that layer and age are shown. **(B)** View in tangential plane of the intracortical axonal arbors of 10 L6CThNs in layer 6. **(C)** Summary data for the total axon length (left) and the number of branch points (right) of the intracortical axons of L6CThNs in L6 across the four age groups (p < 10^-4^ for both metrics, Kruskal-Wallis tests; P4: n = 10 cells, N = 3 animals; P7: n = 14 cells, N = 5 animals; P10: n = 12 cells, N = 5 animals; and P14: n = 12 cells, N = 8 animals). **(D)** As in B but for layer 5 (L5). **(E)** Summary data as in C but for L5 (p < 0.002 for both metrics, Kruskal-Wallis tests). **(F)** As in B but for layer 4 (L4). **(G)** Summary data as in C but for L4 (length: p < 0.003, branch points: p < 0.012, Kruskal-Wallis tests). C, E, G: Pairwise Wilcoxon rank-sum tests with a Bonferroni correction for multiple comparisons were used to compare between consecutive age groups. *, p < 0.05; **, p < 0.01; ***, p < 0.001. Data in C, E, G shown as mean ± SEM. Scale bars: B, D, E: 200 μm.

### Developmental timing differs between dendritic and axonal arbors of layer 6 corticothalamic neurons

Next, we determined whether L6CThN dendrite elaboration followed a similar developmental time course as the intracortical axons of L6CThNs. The apical dendrites of all reconstructed L6CThNs terminated largely in L4 (Figure S2A, S2B). Thus, the apical dendrites of L6CThNs reached their mature vertical extent at the earliest timepoint assessed, P4, in contrast to L6CThN axons which elaborate in L4 late in the second postnatal week. In addition, although the total dendritic length and number of dendritic branches increased between P4-P7 and P10-P14, we observed a decrease in total dendritic branch number between P7-P10 driven primarily by a decrease in branches emerging from the apical dendrite (Figure S2C-E). This result contrasts with the results for axons, which showed no evidence of net pruning. Furthermore, when we compared layer-specific dendritic growth in L4, L5, and L6, we found that significant dendritic growth occurs between P4-P7 across all layers, in contrast to axons, which elaborated primarily in L6 before P7 (Figure S2F-K). These results highlight distinct temporal growth patterns for L6CThN dendrites and axons: L6CThN apical dendrites, which extend only to the middle layers of the cortex, are established across L4-L6 before P4, in contrast to L6CThN intracortical axons which extend to L4 during the second postnatal week.

### Decreasing the excitability of layer 6 corticothalamic neurons increases axon elaboration selectively in layer 6

Prior studies show that neuronal activity affects the growth, branching, and targeting of long-range axonal projections, promoting, for example, the elaboration of corticocallosal and thalamocortical axons^53–58,72^. However, the effects of intrinsic neuronal activity on excitatory intracortical axon development are less clear^49^. To test the effects of reducing L6CThN excitability on the development of their intracortical axons, we selectively expressed the potassium channel Kir2.1 in L6CThNs. We confirmed that Kir2.1 expression reduced the excitability of L6CThNs by comparing the intrinsic properties of L6CThNs in acute slices of somatosensory cortex from P6-P7 mice electroporated with the *Kir2.1* construct or with a control *EGFP* construct. We found that Kir2.1-expressing L6CThNs were significantly more hyperpolarized than control neurons expressing EGFP (Figure S3A), and that the input resistances of Kir2.1-expressing L6CThNs were significantly lower than those of control L6CThNs (Figure S3B), both parameters demonstrating decreased excitability of L6CThNs expressing Kir2.1. To determine whether the decreased excitability of L6CThNs affected axonal development, we compared L6CThN intracortical axons expressing Kir2.1 with control L6CThNs expressing EGFP in P14 mice, a time when L6CThN axons are elaborated in both L6 and L4 (Figure 3A). Decreasing the excitability of L6CThNs resulted in a significant increase in the total number of intracortical axon branches of L6CThNs, although the total intracortical axon length did not significantly differ between Kir2.1 and control L6CThNs (Figure 3B). Given the layer-specific timing of L6CThN intracortical axon development, we next compared the effects of decreased L6CThN excitability on axon arbors in each cortical layer (Figure 3C-H). Reducing L6CThN excitability resulted in a significant increase in both axon length and total number of axonal branch points in L6 (Figure 3F), which also resulted in a significantly broader tangential spread of axons in L6 (convex hull area of L6 axons: WT 27,364.7 ± 4,467.29 μm^2^; Kir2.1: 82,272.1 ± 10,223.7 μm^2^; p < 0.001, Wilcoxon rank-sum test). However, neither total axon length nor the total number of axonal branch points in L4 and L5 were significantly affected by reducing L6CThN excitability (Figure 3G-H). Together, these results indicate that the intracortical axon growth of L6CThNs is negatively regulated by intrinsic neuronal excitability, and that this specifically affects L6 intralaminar axons.

**Figure 3.**
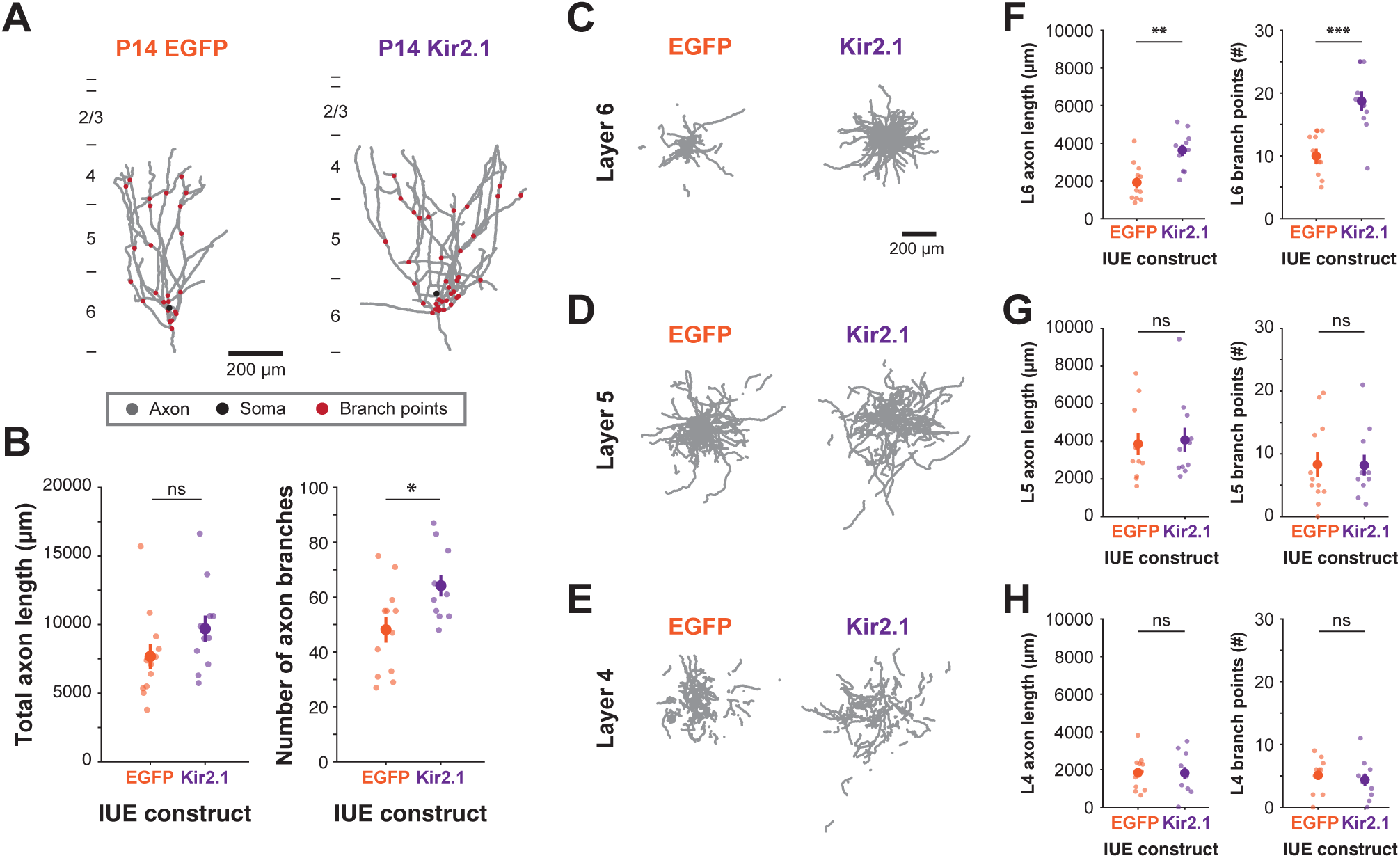
Decreasing the excitability of layer 6 corticothalamic neurons (L6CThNs) via Kir2.1 overexpression results in increased axon elaboration in layer 6 (L6). **(A)** Example axon reconstructions (gray) of L6CThNs expressing EGFP as a control (left) or Kir2.1 (right) from postnatal day 14 (P14) mice. Red circles represent branch points; black circles represent the soma location. **(B)** Summary data showing the total intracortical axon length (left; p = 0.103, Wilcoxon rank-sum test) or number of branches (right; p < 0.033, Wilcoxon rank-sum test) of L6CThNs following expression of control EGFP or Kir2.1. Control P14 data are replotted from Figures 1C and 1D. Black circles represent the mean; gray circles represent individual cells (P14 control EGFP: n = 12 cells, N = 8 animals; P14 Kir2.1: n = 11 cells, N = 7 animals). **(C-E)** Views in tangential plane of the intracortical axonal arbors of L6CThNs expressing control (left, EGFP) or Kir2.1 (right) constructs in layer 6 (L6; C), layer 5 (L5; D) and layer 4 (L4; E). For each experimental group and layer, the 11 cells with axon lengths in that layer closest to the mean of that layer and group are shown. **(F-H)** Summary data showing the total axonal length (left) and total number of branch points (right) in L6 (F), L5 (G) and L4 (H) in L6CThNs expressing control EGFP or Kir2.1 ((F) length: p < 0.002, branch points: p < 10^-4^; (G) length: p = 0.926, branch points: p = 0.877; (H) length: p = 0.878, branch points: p = 0.351, Wilcoxon rank-sum tests). Data in B, F, G and H shown as mean ± SEM. Control data replotted from Figures 1 and 2. Scale bars: A, C-E: 200 µm.

### Decreasing the excitability of layer 6 corticothalamic neurons selectively increases the elaboration of apical dendrites

Next, we asked whether the development of L6CThN dendrites was altered by reducing the intrinsic excitability of L6CThNs via Kir2.1 expression. Although total dendritic length and branch number were not significantly different between L6CThNs expressing Kir2.1 or control EGFP (Figure S4A-B), the total number of apical branches, but not basal branches, was increased in Kir2.1-expressing neurons (Figure S4C-D). Consistent with this result, both the total dendritic length and the number of dendritic branch points were significantly increased in L4 but not in L5 or L6 (Figure S4E-J). Although the development of both L6CThN axons and dendrites are negatively regulated by intrinsic neuronal activity, these effects differ for L6CThN axonal and dendritic processes. Our results show that axonal development is significantly affected in L6, where intralaminar axons are elaborated during the first postnatal week, while dendritic development is most strongly affected in L4, where the apical tufts of L6CThNs in somatosensory cortex are primarily located.

### Layer 6 corticothalamic neurons form synapses preferentially onto parvalbumin-positive inhibitory interneurons in layer 6

We next asked how synapse formation by L6CThN axons proceeded across the two phases of L6CThN development in L6 and L4, a time period when synaptogenesis proceeds rapidly across the cortex^39–42^. In both L6 and L4 in adult mice, L6CThNs preferentially synapse onto PV interneurons relative to more common excitatory neurons, including other L6CThNs^9,16,20–31^. Does preferential synaptogenesis or promiscuous synapse formation followed by selective pruning generate this adult pattern of synaptic connectivity? To test for these two possibilities, we first compared the connection probability for L6CThN→L6 PV interneuron connections with those of L6CThN→L6CThN connections during the first two postnatal weeks using paired whole-cell current-clamp recordings in acute slices of S1 from *Ntsr1-Cre;tdTomato;G42* mice (Figure 4A-D). Few L6CThN→L6 PV interneuron or L6CThN→L6CThN unitary synaptic connections were detected during the first postnatal week (Figure 4E, Table S1). After P8, the probability of detecting L6CThN→L6 PV interneuron unitary synaptic connections increased significantly, reaching 35% in the P9-P15 age range. In contrast, the probability of detecting L6CThN→L6CThN connections remained low, averaging 1.4% from P9 to P15 (Figure 4E, 4F). Indeed, the probability of identifying a L6CThN→L6CThN unitary synaptic connection from P9-P15 was not significantly different from the L6CThN→L6CThN connection probability at P3-P8 (Figure 4F, Table S1). In addition, we found that the amplitude of the first unitary excitatory postsynaptic potential (uEPSP) elicited was significantly greater for detected L6CThN→L6 PV connections than for unitary synaptic connections between L6CThNs (Figure 4G, 4H). Interestingly, although L6CThNs form facilitating intracortical synapses in older animals^16,20,21,23,24,31,73^, on average, both the L6CThN→L6 PV interneuron synapses and the few L6CThN→L6CThN connections detected did not exhibit facilitating paired-pulse ratios during the first two postnatal weeks (Figure 4I, 4J). Therefore, as electrically detectable synapses increase during the first two postnatal weeks, not only do tested L6CThN→L6 PV connections have a higher connection probability than L6CThN→L6CThN pairs, they also exhibit a higher overall connection strength than L6CThN→L6CThN connections. Together, these results using paired recordings show that L6CThNs preferentially form functional synaptic connections onto L6 PV interneurons during the period of initial synaptogenesis.

**Figure 4.**
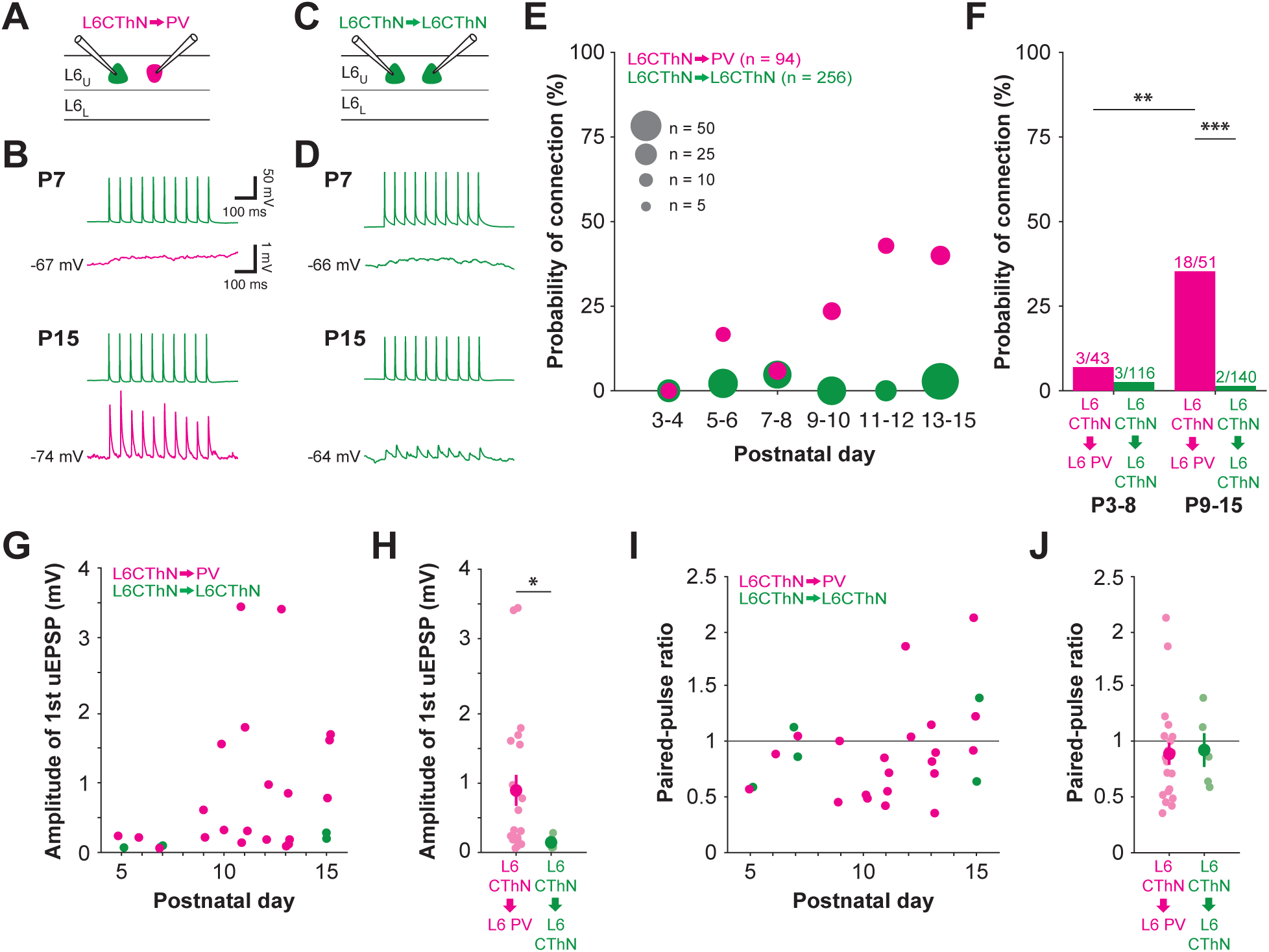
Layer 6 corticothalamic neurons (L6CThNs) form synapses preferentially onto layer 6 parvalbumin (L6 PV) inhibitory interneurons. **(A)** Experimental configuration showing whole-cell patch clamp recordings from pairs composed of a L6CThN and a L6 PV interneuron. **(B)** Unitary L6CThN→L6 PV interneuron synaptic connections recorded in acute slices of somatosensory cortex (S1) from a postnatal day 7 (P7, top) and P15 (bottom) mouse. Action potentials initiated in a presynaptic L6CThN (top, single trace, 20 Hz) and the average unitary excitatory postsynaptic potentials (uEPSPs) recorded in a L6 PV interneuron (lower trace) are shown. **(C)** Experimental configuration as in A but for two L6CThNs. **(D)** Unitary L6CThN→L6CThN synaptic connections recorded in acute slices of S1 from a P7 (top) and P15 (bottom) mouse shown as in B. **(E)** Probability of connection for tested L6CThN→L6 PV (magenta) and L6CThN→L6CThN (green) potential connections from P3 to P15. Symbol diameters represent the number of potential connections tested for each time bin (L6CThN→L6 PV: n = 94 tested potential connections, N = 43 animals; L6CThN→L6CThN: n = 256 tested potential connections from 128 pairs, N = 63 animals, see Table S1). **(F)** Probability of connection for tested L6CThN→L6 PV and L6CThN→L6CThN potential connections in two age ranges, P3-P8 (L6CThN→L6 PV: N = 29 animals; L6CThN→L6CThN: N = 20 animals) and P9-P15 (L6CThN→L6 PV: N = 23 animals; L6CThN→L6CThN: N = 34 animals). Chi-squared test of independence was performed to test for significant differences in cell-type-specific connectivity (p < 10^-13^). Fisher’s exact tests with Bonferroni corrections for multiple comparisons were used for pairwise comparisons between groups. **(G)** Average amplitudes of the first uEPSP for each detected L6CThN→L6 PV and L6CThN→L6CThN connection plotted across age. Statistical significance of changes in EPSC amplitude were assessed using permutation tests of the Spearman coefficient of correlation (p = 0.0794 for n = 21 L6CThN→L6 PV connections, N = 16 animals; p < 10^-4^ for n = 5 L6CThN→L6CThN connections, N = 3 animals). **(H)** Summary data comparing the average amplitudes of the first uEPSPs for detected L6CThN→L6 PV and L6CThN→L6CThN connections (L6CThN→L6 PV: 0.90 ± 0.22 mV; L6CThN→L6CThN: 0.15 ± 0.042 mV, p < 0.032, Wilcoxon rank-sum test). **(I)** Paired-pulse ratio (PPR) for detected L6CThN→L6 PV and L6CThN→L6CThN connections. Statistical significance of changes in PPR were assessed using permutation tests of the Spearman coefficient of correlation (p = 0.0996 for L6CThN→L6 PV; p = 0.3613 for L6CThN→L6CThN). **(J)** Summary data comparing the average PPR for detected L6CThN→L6 PV and L6CThN→L6CThN connections (p = 0.6027, Wilcoxon rank-sum test). *, p < 0.05; **, p < 0.01; ***, p < 0.001. Data in H and J shown as mean ± SEM.

### Layer 6 corticothalamic neurons preferentially activate layer 4 parvalbumin-positive inhibitory interneurons during synapse formation in the second postnatal week

As L6CThNs also preferentially provide input to PV interneurons in L4 in older mice^20^, we tested whether preferential synaptogenesis also occurs for these interlaminar L6CThN connections, comparing three time periods: 1) P7-P8, when axon elaboration has largely taken place in L6 but not in L4; 2) P10-P11, following the pause in axon development; and 3) P14-P15, following the growth of L6CThN axons in L4. We opted to use channelrhodopsin-2-assisted circuit mapping (CRACM^74^) as recording unitary synaptic connections between L6 and L4 in brain slices is particularly challenging. To confirm whether ChR2 effectively activates L6CThNs across these ages, we recorded first from L6CThNs in brain slices from mice hemizygous or homozygous for *ChR2-EYFP*. L6CThNs were reliably activated in mice homozygous, but not hemizygous, for *ChR2-EYFP* at all three ages (Figure S5A-E). Therefore, we used acute slices of barrel cortex from *Ntsr1-Cre;ChR2-EYFP+/+;G42* mice for comparing L6CThN input to L4 PV interneurons and L4 excitatory cells. We optogenetically activated ChR2-expressing L6CThN axons with brief flashes of light while recording responses simultaneously from the two postsynaptic cell types (Figure 5A, 5B). At P7-P8, responses recorded simultaneously from pairs composed of a L4 PV interneuron and a L4 excitatory cell were weak or absent following L6CThN optogenetic activation (Figure 5C). At P10-P11, we detected weak responses to L6CThN activation in many L4 PV interneurons and some L4 excitatory neurons. The response amplitudes recorded from L4 PV interneurons were significantly greater than those recorded simultaneously from L4 excitatory neurons (Figure 5D). Even at this early developmental stage, the responses of some L4 excitatory neurons were dominated by previously reported disynaptic inhibitory responses following L6CThN activation^20,25^. At P14-P15, depolarizing responses recorded from L4 PV neurons were significantly larger than those from L4 excitatory neurons (Figure 5E). Notably, the responses of L4 excitatory neurons at P14-P15 were all hyperpolarizing, reflecting disynaptic inhibition elicited by photostimulation of L6CThNs. These results show that L6CThNs preferentially activate L4 PV interneurons beginning with initial formation of their synapses. This result is consistent with the preferential formation of functional, electrically detectable synaptic connections by L6CThNs onto PV interneurons both locally within L6 and in L4.

**Figure 5.**
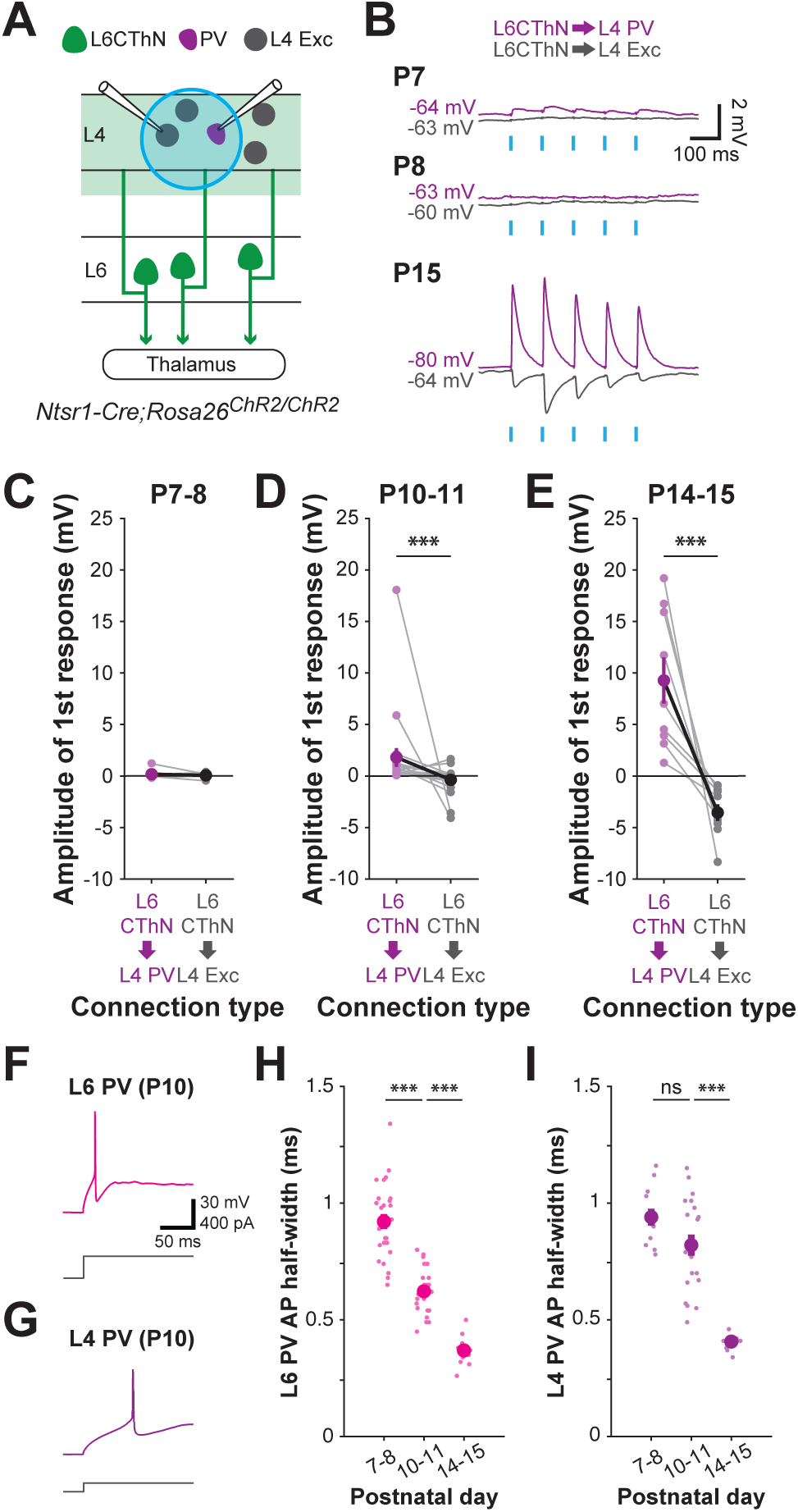
Layer 6 corticothalamic neurons (L6CThNs) preferentially activate layer 4 (L4) parvalbumin (PV) inhibitory interneurons during synapse formation in the second postnatal week. **(A)** Experimental configuration showing whole-cell recordings from pairs composed of a L4 PV (magenta) and a L4 excitatory (green) neuron while photostimulating channelrhodopsin2-expressing L6CThN axons in acute slices of somatosensory cortex from *Ntsr1-Cre;Rosa26^ChR^*^2^*^/ChR^*^2^ mice. **(B)** Example average postsynaptic responses from pairs of L4 neurons (PV: magenta; excitatory: green) in response to optogenetic stimulation of L6CThN axons in slices from postnatal day 7 (P7; top), P8 (middle), and P15 (bottom) mice. Cyan bars indicate light stimulation (five 3 ms light pulses at 10 Hz). **(C)** Summary data showing the average amplitudes of the postsynaptic responses to the first optogenetic stimulation for L4 PV and excitatory neurons recorded at P7-P8. Gray lines connect pairs recorded simultaneously (p = 0.751, Wilcoxon signed-rank test, n = 12 pairs, N = 4 animals). **(D)** As in C, but for pairs recorded at P10-11 (p < 0.001, Wilcoxon signed-rank test, n = 20 pairs, N = 5 animals). **(E)** As in C, but for pairs recorded at P14-15 (p < 0.001, Wilcoxon signed-rank test, n = 9 pairs, N = 4 animals). **(F)** Example response to a current pulse near rheobase of a L6 PV interneuron (top) to the onset of a depolarizing current pulse (bottom) at P10. **(G)** As in G, but for a L4 PV interneuron at P10. **(H)** Width at half-maximum of the action potential (AP half-width) of L6 PV interneurons at P7-8, P10-11, and P14-15. Large circles represent group means; small circles represent individual cells (p < 10^-10^, Kruskal-Wallis test for change between ages; P7-P8: n = 26 cells, N = 12 animals; P10-P11: n = 22 cells, N = 9 animals; P14-P15: n = 13 cells, N = 5 animals). **(I)** AP half-widths for L4 PV interneurons at P7-8, P10-11, and P14-15 (p < 10^-5^, Kruskal-Wallis test for change between ages; P7-P8: n = 12 cells, N = 4 animals; P10-P11: n = 20 cells, N = 5 animals; P14-P15: n = 10 cells, N = 4 animals). H, I: Pairwise Wilcoxon rank-sum tests with a Bonferroni correction for multiple comparisons were used to compare between consecutive age groups. *, p < 0.05; **, p < 0.01; ***, p < 0.001. Data in C, D, E, G and H shown as mean ± SEM.

### The maturation of parvalbumin inhibitory interneuron action potential properties begins earlier in layer 6 relative to layer 4

Based on the two phases of axonal growth of L6CThNs, first in L6 and then in L4, we predicted that the development of L6CThN→L6 PV interneuron synapses precedes the development of L6CThN→L4 PV synapses. The results from our paired recordings in L6 showed that the connectivity rate for unitary L6CThN→L6 PV interneuron synapses is similar to more mature animals by P9 (37% in P13-P47 mice^16^), a developmental time when L6CThN axons have elaborated in L6. In addition, the larger uEPSPs we recorded in L6 PV interneurons, those with amplitudes greater than 0.5 mV, were detected as early as P9 (Figure 4G). Although using a different approach, our ChR2-circuit mapping suggested a later increase in the number and/or the strength of L6CThN→L4 PV interneuron synapses; L4 PV responses were significantly larger at P14-P15 (9.3 ± 2.2 mV, mean ± SEM) than at P10-P11 (1.81 ± 0.90 mV; p < 0.0004, Wilcoxon rank-sum test; Figure 5D, 5E), correlating with the period of L6CThN axon elaboration in L4 between P10-P11 and P14-P15 (Figure 2F, 2G). Since the maturation of PV interneurons is influenced by excitatory synaptic input^75^, we hypothesized that a difference in the timing of L6CThN axon formation and synaptogenesis in L6 and L4 may be correlated with differences in the timing of PV interneuron maturation in these two cortical layers. We tested this possibility by comparing the intrinsic electrophysiological properties of L6 and L4 PV interneurons at three developmental timepoints, P7-P8, P10-P11, and P14-P15. We found that the development of the canonical narrow-spiking phenotype^76–79^ began earlier in L6 PV interneurons than in L4 PV interneurons (Figure 5F-I, S5F-I). For example, the average action potential (AP) half-width of L6 PV interneurons decreased significantly between P7-8 and P10-11 and also between P10-11 and P14-P15 while, for L4 PV interneurons, it significantly decreased only during the later time period (Figure 5H, 5I). Similarly, changes in the maximum rate of rise and fall of the APs occurred during both the earlier and later time periods for L6 PV interneurons, but only during the later time period for L4 PV interneurons (Figure S5G, S5I). These results show that narrow-spiking properties develop later in L4 PV interneurons than in L6 PV interneurons. Taken together, our results show that the maturation of intrinsic electrophysiological properties of L4 and L6 PV interneurons is correlated with the development of L6CThN axon morphology and preferential synaptic connectivity onto PV interneurons in those layers.

### Layer 6 corticothalamic neurons do not form silent synapses during the period of peak neonatal synaptogenesis in layer 6

Our results indicate that L6CThNs preferentially form functional excitatory synapses onto PV interneurons over neighboring excitatory neurons, as assessed using current-clamp recordings. However, silent synapses, which contain NMDA receptors but not AMPA receptors and are thus not typically detected in current clamp recordings at physiological resting membrane potentials, have been observed in neonatal cortical circuits, including in L6^63,64,80–87^. Thus, it is possible that L6CThNs initially form promiscuous silent synapses onto both L6CThNs and PV interneurons, after which preferential unsilencing occurs at L6CThN→PV synapses via selective insertion of AMPA receptors. To test this possibility, we compared the probability of connection for L6CThN→L6CThN and L6CThN→L6 PV interneuron pairs while candidate postsynaptic neurons were voltage-clamped alternately at membrane holding potentials of +40 mV and -85 mV to detect NMDA-mediated as well as AMPA-mediated synaptic currents (Figure 6). We focused on slices from P6-P9 mice since this age range coincides both with the ages silent synapses have been reported in L6^63,64^ and with the time period of increasing L6CThN→L6 PV connectivity measured in our current-clamp experiments (Figure 4E). We did not observe any silent synapses, defined as connections detected at a membrane holding potential of +40 mV but not at -85 mV, nor any AMPA-mediated connections, in 42 potential L6CThN→L6CThN connections that were tested (Figure 6A, 6B). We also did not observe any L6CThN→PV silent synapses among the 15 potential L6CThN→L6 PV interneuron connections tested (Figure 6C, 6D), although we did detect three L6CThN→PV connections at -85 mV (20%), one of which also had detectable uEPSCs at +40 mV (Figure 6C). These results show that L6CThNs do not form silent synapses at P6-P9 and argue against a role for promiscuous silent synapse formation and selective unsilencing in L6CThN preferential synapse formation.

**Figure 6.**
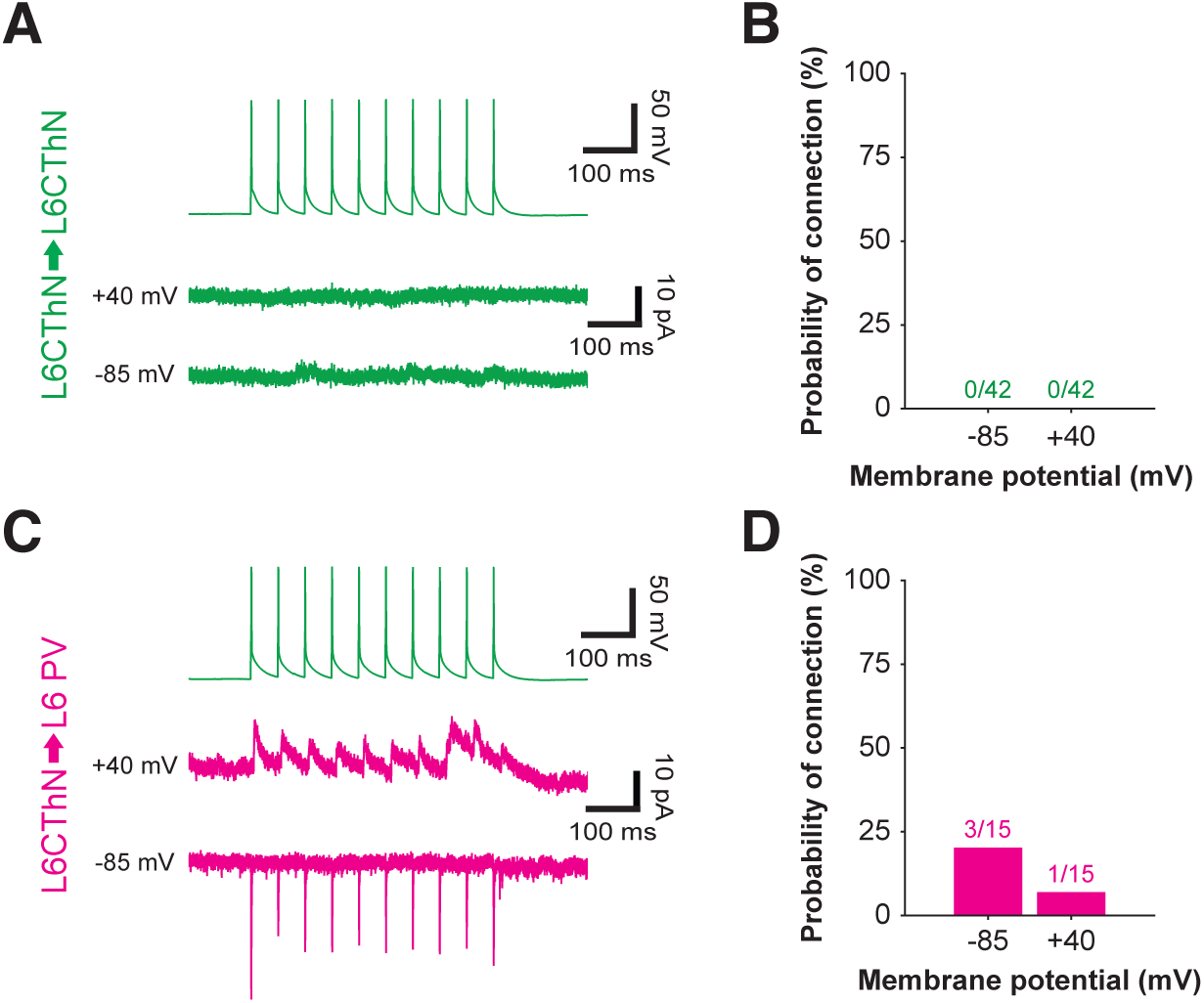
Layer 6 corticothalamic neurons (L6CThNs) do not form silent synapses during synaptogenesis in layer 6. **(A)** Example average traces recorded in a L6CThN neuron held at -85 mV (bottom trace) and +40 mV (middle trace) in response to action potentials evoked in a presynaptic L6CThN (20 Hz, top trace, green). **(B)** Probability of detecting a L6CThN→L6CThN connection at -85 mV and +40 mV membrane potentials at postnatal day 6 (P6) to P9 (n = 42 connections tested from 21 L6CThN-L6CThN pairs, N = 21 animals). **(C)** Example traces as in A, but for a tested L6CThN→L6 PV connection. **(D)** As in B, but for tested L6CThN→L6 PV connections. The connection identified at +40 mV was also detected when measured at -85 mV (n = 15 tested connections, N = 9 animals).

### Reducing calcium-permeable AMPA receptors does not affect the development of layer 6 corticothalamic neuron synapses onto inhibitory interneurons

PV and somatostatin-positive (SOM) interneurons, both of which are derived developmentally from the medial ganglionic eminence^88–90^, express a high proportion of calcium-permeable AMPA receptors^91–94^. These receptors have been shown to mediate activity-dependent synaptic plasticity and long-term potentiation^91,95–99^, raising the possibility that they could contribute to L6CThN→PV synapse formation. In our experiments testing L6CThN→L6 PV connectivity in voltage clamp, the three L6CThN→L6 PV connections we detected showed inward rectification of synaptic currents, characteristic of calcium-permeable AMPA receptors^100–102^. To test if reducing the calcium permeability of AMPA receptors in PV interneuron synapses alters the development of L6CThN→L6 PV connectivity, we overexpressed the Q/R-edited calcium-impermeable AMPA receptor subunit GluA2 in PV interneurons. Since PV expression starts late in the second postnatal week^103^, we used an *Lhx6-CreER^T^*^2^ mouse line, in which CreER^T2^ is expressed in both PV and SOM interneurons^104^, crossed to a *LSL-EGFP-Gria2* mouse line^91,105^, to selectively overexpress GluA2 in Lhx6 interneurons starting in the first postnatal week. We confirmed that GluA2 overexpression following postnatal tamoxifen injections (P1-P3) altered the calcium permeability of AMPA receptors of Lhx6 interneurons, observing greater inward rectification of synaptic currents (Figure S6A-H), without affecting their intrinsic electrophysiological properties (Figure S6I) or the total number or distribution of PV or SOM neurons (Figure S6J-L).

To test if reduced calcium permeability of AMPA receptors altered L6CThN→L6 Lhx6 interneuron synapse formation, we recorded from neuron pairs composed of an Lhx6 interneuron in L6 and a L6CThN retrogradely labeled from VPM in S1 slices from P14-P24 *Lhx6-CreER;EGFP-Gria2* and control *Lhx6-CreER;EYFP* mice (Figure S7A-C). We found that the probability of connection, average amplitude, and paired-pulse ratio of L6CThN→L6 Lhx6 connections did not significantly differ between slices from control *Lhx6-CreER;EYFP* and *Lhx6-CreER;EGFP-Gria2* mice (Figure S7D-F), indicating that overexpressing GluA2 receptors during the first two postnatal weeks does not affect the probability of connection or the connection strength of L6CThN→L6 Lhx6 interneuron synapses at P14-P24. Although calcium-permeable AMPA receptors have been implicated in synaptic plasticity of excitatory synapses onto inhibitory interneurons in other contexts^91,97–99^, our results support the hypothesis that the formation of L6CThN→L6 PV interneuron synapses does not require calcium-permeable AMPA receptors.

### Development of layer 6 parvalbumin inhibitory interneuron synapses onto layer 6 corticothalamic neurons

Not only do L6CThNs preferentially synapse onto PV interneurons, but L6CThNs also receive substantial input from PV interneurons^9,16,20–26^. Whether L6CThN→L6 PV interneuron synapses and L6 PV interneuron→L6CThN synapses are established in parallel, as suggested by some studies of connectivity between inhibitory and excitatory neurons in neonates^79^, or with a temporal delay, as suggested by other work^106^, is not known. In our whole-cell recordings from L6CThN-L6 PV interneuron pairs, we detected rare unitary L6 PV interneuron→L6CThN connections as early as P7-8 (Figure 7A-D, Table S2). We found that the probability of connection increased rapidly after P8 (Figure 7D, 7E; P9-P15: 61%, n = 31 of 51 tested connections, N = 23 mice; Fisher’s exact test, p < 10^-6^), reaching a level similar to that reported for older animals (54%)^16^.This time course is similar to the time course observed for the development of L6CThN→L6 PV interneuron connection probability (Figure 4E, 4F; Table S1). In addition, we found that many of the L6 PV interneuron-L6CThN pairs we tested formed reciprocal synaptic connections during the first two postnatal weeks. Of the neuron pairs with identified L6CThN→L6 PV interneuron connections recorded in slices from P9-P15 mice, 89% formed a reciprocal L6 PV interneuron→L6CThN connection (Figure 7F). For the L6 PV interneuron→L6CThN connections detected, 52% were reciprocal, similar to what has been described in older animals (43%)^16^. Reciprocal connectivity was significantly higher than expected if L6CThN→L6 PV interneuron and L6 PV interneuron→L6CThN connections formed independently (p < 0.0024, Chi-square test of independence). These results indicate that L6CThN→L6 PV interneuron and L6 PV interneuron→L6CThN connections develop over similar time courses, with a bias in favor of forming reciprocal connections.

**Figure 7.**
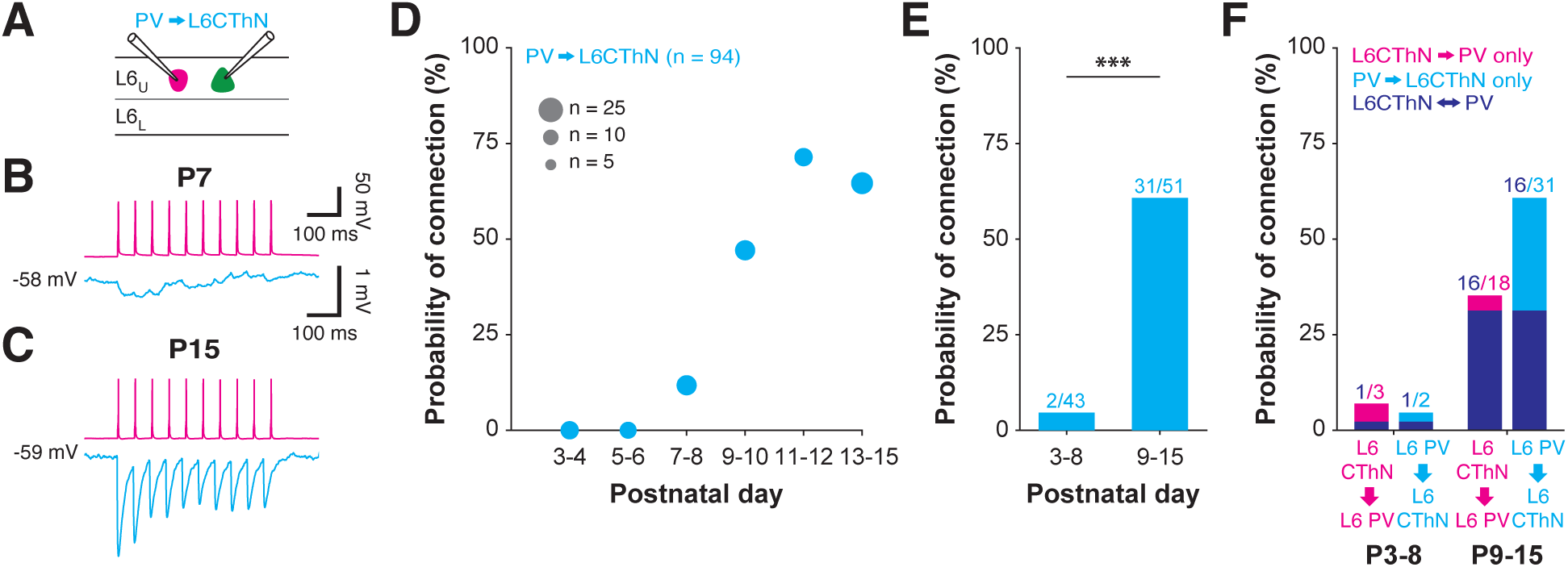
Maturation of the synaptic connectivity of layer 6 parvalbumin (L6 PV) inhibitory interneurons. **(A)** Experimental configuration showing whole-cell patch clamp recordings from pairs composed of a L6 PV interneuron and a L6CThN. **(B-C)** Unitary L6 PV→L6CThN synaptic connections recorded in acute slices of somatosensory cortex from a postnatal day 7 (P7; B) and P15 (C) mouse. Action potentials initiated in a presynaptic L6 PV interneuron (above, single trace, 20 Hz) and the average unitary inhibitory postsynaptic potentials (uIPSPs) recorded in a L6CThN interneuron (lower trace) are shown. The P7 L6 PV→L6CThN connection shown in B and the P15 L6 PV→L6CThN connection shown in C are from the same pairs displayed in Figure 1B. **(D)** Probability of connection for tested L6 PV→L6CThN connections from P3 to P15. Symbol diameters represent the number of potential connections tested for each time bin (n = 94 tested connections, N = 43 mice; see Table S2). **(E)** Probability of connection for tested L6 PV→L6CThN potential connections from P3-P8 (N = 20 animals) and from P9-P15 (N = 23 animals; p < 10^-6^, Fisher’s exact test). **(F)** Probability of connection for unidirectional L6CThN→L6 PV (magenta), unidirectional L6 PV→L6CThN (cyan), and reciprocally connected (blue) L6CThN-L6 PV tested pairs from P3-8 and from P9-P15.

## DISCUSSION

We found that the intracortical axons of L6CThNs exhibited two periods of growth. L6CThN intracortical axons extended within L6 before P7, then paused before exhibiting translaminar growth to L4 between P10 and P14. Decreasing the excitability of L6CThNs via overexpression of Kir2.1 selectively increased the outgrowth and branch elaboration of their intralaminar axons, indicating that intrinsic neuronal activity negatively regulates the earlier, intralaminar phase of growth. In contrast, the dendrites of L6CThNs reached their vertical extent prior to P4, and intrinsic neuronal activity negatively regulated their apical dendritic tuft branching in L4. Next, we showed that the preference of L6CThNs for forming functional, AMPA-receptor-containing synaptic connections onto PV interneurons in both L6 and L4 of adult mice emerges during initial synapse formation, rather than resulting from promiscuous synapse formation followed by selective pruning of inappropriate synapses. We did not detect L6CThN silent synapses during early synaptogenesis, further arguing against the indiscriminate formation of silent synapses by L6CThNs followed by cell-type-specific synapse unsilencing. Together, these results indicate that preferential synaptogenesis underlies the adult patterns of synaptic connectivity between L6CThNs and PV interneurons.

By reconstructing the intracortical axons of filled L6CThNs at four time points during the first two postnatal weeks, we found that they first elaborated locally within L6 and then, after a pause, translaminarly to L4. Prior studies of intracortical pyramidal neuron axon development in other cortical layers in rodents (L2/3, L4 and L5)^46,48–51^, and of L6 pyramidal neurons in the visual cortex of ferrets ^47^, have not reported a similar pause in axonal growth across cortical layers. This difference may be due to L6CThN-specific spatiotemporal patterns of axon development in rodents. Alternatively, it may reflect differences in experimental approach, since here we focused on a well-defined cell type and assessed intact axonal arbors in cleared brains from mice P4 to P14. Similar to prior studies in rodents^46,48–51^, we did not detect net overgrowth of intracortical axons followed by pruning. However, dynamic growth and elimination of axon branches and protrusions that do not lead to net changes in axon metrics may still occur^49^.

In addition to identifying layer-specific differences in the developmental timing of L6CThN axon intracortical elaboration, we also found that neuronal activity affects L6CThN axon development differently across layers. Reducing the intrinsic neuronal activity of L6CThNs via Kir2.1 overexpression increased length and branching of L6CThN axons in L6, but not in L5 or L4, indicating that neuronal activity negatively regulates the early phase of axon growth and elaboration specifically within L6. In contrast, prior studies have shown that reducing neuronal excitability via Kir2.1 expression results in reduced axon length and branching in several contexts, including the axons of certain cortical inhibitory neuron subtypes^107^, long-range callosal axons of L2/3 excitatory neurons^53–58^, and thalamocortical axons *in vitro* and *in vivo*^72,108,109^. These studies show a positive effect of intrinsic neuronal activity on axon elaboration in these cases. Although the negative regulation of axon length and branch number we observed here has not been previously described for cortical neurons, negative regulation by intrinsic neuronal activity has been proposed to contribute to refining other long-range axonal projections, such as those from the olfactory epithelium to the olfactory bulb and from retinal ganglion cells to their central targets^110–113^. Furthermore, reducing intrinsic neuronal activity in L2/3 excitatory neurons leads to an increase in the number of transient short intracortical axonal protrusions^49^. Therefore, intrinsic activity of L6CThNs may specifically inhibit the initiation, stabilization, or elongation of L6CThN axonal protrusions during development.

The development of L6CThN dendrites differed from that of intracortical axons in both timing and activity dependence. In contrast to L6CThN axons, for which most of the growth and branching in L4 was delayed until P10-P14, the apical dendrites of L6CThNs extended into L4 by P4, in agreement with prior work^114^. Since the apical dendrites of reconstructed L6CThNs in S1 did not extend beyond L4, as observed in rats^19^, our results are consistent with studies of L5 pyramidal neurons showing that their layer 1 apical dendrites reach their full pial extent in the first postnatal days before subsequently elaborating additional tuft, oblique and basal dendrites^115–119^. We found that intrinsic neuronal activity negatively regulates L6CThN apical dendrite complexity, in contrast to the effects on supragranular cortical pyramids, for which decreasing their intrinsic excitability inhibited dendritic development^59,120^. Furthermore, decreasing the intrinsic excitability of L6CThNs affected dendritic growth and branching specifically in apical dendrites of L4, in contrast to the intracortical axons of L6CThNs for which intralaminar growth within L6 was specifically affected. Although the molecular mechanisms linking the intrinsic activity of L6CThNs to the elaboration of their intralaminar axons and their apical dendrites remain unclear, L6CThN activity may act via activity-dependent transcription factors, as has been demonstrated in cortical inhibitory subtypes^107^, or by regulating the expression of guidance molecules as for thalamocortical axons^72^. Our results are consistent with a more prominent role for intrinsic neural activity in regulating early dendritic and axonal development during the first postnatal week, when L6CThN intralaminar axons and apical dendrites in L4 develop. These may represent temporally specific effects, as has been shown for cortical inhibitory neurons^107^.

Our results show that the timing of the formation of L6CThN synapses onto L6 PV interneurons, and the formation of L6 PV interneuron synapses onto L6CThNs, was similar, which may be a shared property of pyramid-PV connections across layers^79^ (but see ^106^). The development of L6CThN synaptic connectivity broadly coincided with maturation of feedforward inhibitory circuits mediated by PV interneurons at the beginning of the second postnatal week^90,121–123^. Our results suggest that the timing of L6CThN synaptogenesis differs between cortical layers, with synapse formation in L6 preceding synapse formation in L4. Prior studies demonstrate that the electrophysiological properties of PV interneurons exhibit progressive maturation across the age range we tested^76–79^. We found that the maturation of PV interneuron action potential properties in L4 was delayed relative to L6 PV interneurons, and coincided with a fivefold increase in the strength of L6CThN input to L4 PV interneurons. Since the maturation of PV interneurons is influenced by excitatory synaptic input^75^, our results raise the possibility that a difference in the timing of L6CThN synaptogenesis in L6 and L4 contributes to these differences in PV interneuron maturation in the two layers. Together, these results suggest that, even for a single cell type such as L6CThNs, the timing of developmental events like synaptogenesis differs across cortical layers and correlates with layer-specific development of synaptic partners.

Both promiscuous synapse formation followed by pruning and selective synaptogenesis have been described for cortical inhibitory neurons^1^. For example, recent electron microscopic analyses showed that cortical inhibitory neurons that preferentially target the somas and axon initial segments of pyramidal neurons prune off-target synapses to increase their specificity for these subcellular compartments during development^40^ (but see ^60^). In contrast, cortical inhibitory neurons that synapse onto apical dendrites exhibit preferential synaptogenesis^40^. Similarly, preferential synaptogenesis contributes to the development of immunohistochemically identified synapses between basket cell subtypes and different classes of L5 pyramidal neurons^3^. Here, we show that an excitatory pyramidal subtype exhibits preferential functional synaptogenesis during the first two postnatal weeks. L6CThNs formed functional AMPA-containing synapses onto PV interneurons in L4 and L6 while avoiding the more numerous excitatory neurons found in each layer, similar to the selectivity observed in more mature mice^9,16,20–31^. In addition, we tested the possibility that this preferential synapse formation was generated following the promiscuous formation of silent synapses, with selective unsilencing of L6CThN→L6 PV interneuron synapses via activity-dependent AMPA receptor insertion, another potential mechanism for generating preferential connectivity for excitatory cortical cell types. We did not detect silent synapses during the peak period of L6CThN synaptogenesis, arguing against this mechanism. Our results also indicate that the silent synapses previously reported in neonatal L6^63,64^ must represent synapses formed by cell types other than L6CThNs, including possibly corticocortical or thalamocortical projection neurons. Calcium-permeable AMPARs are highly expressed in PV interneurons^91–94^, and they contribute to synaptic plasticity of excitatory synapses onto inhibitory interneurons in older animals^91,95,97–99^, thus raising the possibility that they are required for the development of specific inputs to PV interneurons. However, we show that they are not required for forming L6CThN synapses onto PV interneurons. Together, our results show that selective synaptogenesis forms a microcircuit in which L6CThNs bias their outputs to PV inhibitory neurons, an important inhibitory pathway implicated in regulating cortical sensory responses^25–27,34–38^.

Whether the preferential synaptogenesis observed for L6CThN connections extends to other excitatory cell types within the cortex, or whether both selective synaptogenesis and pruning of off-target synapses contribute to generating the cell-type-specific patterns of excitatory cortical connectivity, remains to be tested. Furthermore, comprehensively addressing these possibilities will require anatomical as well as functional assessments of synapse formation. In addition, the precise molecular mechanisms by which L6CThNs preferentially form synapses onto PV neurons while avoiding other pyramidal cell dendrites during development remain unclear, as do potential consequences of perturbing such selectivity on cortical circuit function. Overall, our results show that cell-type-biased synaptogenesis underlies the formation of functional cell-type-specific L6CThN synaptic connections and represent important steps in understanding the formation of cell-type-biased excitatory connectivity in the neocortex.

## Supporting information

GutmanWei, Sudarsanam et al Supplemental Figures and Tables

## ACKNOWLEDGEMENTS

This work was supported by NIH Training Grant T32EY017203 (AYG-W), R25 NS107167 (SMS-K), a National Science Foundation Predoctoral Fellowship (AYG-W), a Kavli Neuroscience Discovery Institute Distinguished Graduate Student Fellowship (SS), R03TR004616 (ALK), P30NS050274 (ALK), Johns Hopkins COVID Bridge Funding (SPB) and The Harry Fred Fund (SPB). We thank Dwight E. Bergles for generously sharing the *ROSA26-CAG-loxP-stop-loxP-EGFP-Gria2-WPRE* mouse, the NINDS Microscopy core, Aleksandr Smirnov, Michelle Pucak, Abby Bush and Jakub Ziak for assistance with clearing and lightsheet imaging and members of the Brown lab for their helpful comments.

## AUTHOR CONTRIBUTIONS

AYG-W, SS, ALK, and SPB conceived of the overall study, approach, and specific experiments. AYG-W and SS performed experiments with assistance from AGC, NS, AS, LGC and SMS-K. AYG-W and SS analyzed the data with assistance from AGC. AA generated the *ROSA26-CAG-loxP-stop-loxP-EGFP-Gria2-WPRE* mouse. AYG-W and SPB wrote the paper with input from all authors.

## MATERIALS AND METHODS

### Mice

All experimental procedures were approved by the Johns Hopkins Animal Care and Use Committee and were conducted in accordance with the guidelines of the National Institutes of Health and the Society for Neuroscience. All mouse lines were maintained on a mixed background composed primarily of C57BL/6J and CD-1 strains. Mice of either sex were used for experiments, and all animals were reared on a 14.5/9.5 hr light/dark cycle and given *ad libitum* access to food and water.

To determine the synaptic connectivity of L6CThNs, a Neurotensin receptor-1 Cre recombinase line (*Ntsr1-Cre*, Gensat 220, MMRC 017266-UCD^68^), which expresses Cre recombinase in L6CThNs in primary sensory cortex^20,26,34^, was used to target L6CThNs for recording, optogenetic stimulation, and *in utero* electroporation. To optogenetically activate L6CThNs, *Ntsr1-Cre* mice were crossed with a mouse line carrying a Cre-dependent *loxP-STOP-loxP-channelrhodopsin-2-EYFP* construct in the Rosa26 locus (Allen Institute *Ai32*, Jackson 024109^129^) and a *Gad1-GFP* transgenic line to fluorescently label a subset of PV interneurons (*G42*, Jackson 007677^127^). To increase the efficiency of optogenetic activation of L6CThNs, mice used for characterizing L6CThN inputs to L4 were homozygous for *ChR2-EYFP*, in addition to being hemizygous for *Ntsr1-Cre* and *G42* (See *Method Details: Optogenetic stimulation of L6CThNs*).

For paired recording experiments, *Ntsr1-Cre* mice were crossed with a Cre-dependent *loxP-STOP-loxP-tdTomato* reporter line (Allen Institute Ai14 or Ai9, Jackson 007905 and 007908^128^) to fluorescently label L6CThNs as well as the *G42* line. Mice used for targeted recordings of both L6CThNs and PV interneurons were hemizygous for *Ntsr1-Cre*, *tdTomato*, and *G42*. All *G42* mice were maintained as heterozygotes.

To determine the role of calcium-permeable AMPA receptors in inhibitory neurons on the formation of L6CThN→PV synapses, a LIM homeobox protein 6 (Lhx6) inducible Cre recombinase knockin line (*Lhx6-CreER*, Jackson 010776^104^) was used to target Lhx6 interneurons, which include PV and somatostatin-positive (SOM) interneurons, for fluorescent labeling and transgenic manipulation. *Lhx6-CreER* mice were crossed with mice carrying a Cre-dependent *loxP-STOP-loxP-EGFP-Gria2* allele to selectively overexpress the GluA2 subunit (*Lhx6-CreER;EGFP-Gria2*^91,105^), or a *loxP-STOP-loxP-EYFP* control allele (*Lhx6-CreER;EYFP*; Allen Institute Ai3, Jackson 007903^128^). All *Lhx6-CreER* mice were maintained and analyzed as heterozygotes, and all experimental mice were hemizygous for either the *EGFP-Gria2* or *EYFP* alleles generated as littermates from crosses of *Lhx6Cre^ERT^*^2^*^/+^* and *Rosa26^EYFP/EGFP-Gria^*^2^ parents.

### In utero electroporation

Homozygous *Ntsr1-Cre* males were crossed to wild-type CD-1 females to generate timed-pregnant female mice. Noon of the day on which a vaginal plug was observed was designated as embryonic day 0.5 (E0.5). *In utero* electroporations were performed on the evening of E11 or the morning of E12 to target L6CThNs for transfection. Pregnant mice were anesthetized with 2% isoflurane and placed on a heating pad. The abdomen was shaved and cleaned, 100 μL of the local anesthetic bupivacaine-hydrochloride (2.5 mg/mL; Sigma B5274-5G) was applied, and a 1.5-2 cm laparotomy was made. Pups were extracted and rinsed with sterile warmed PBS. Pups were injected in one lateral ventricle using glass needles pulled on a vertical pipette puller (Narishige PC-10), and electroporated with five 36 V pulses of 50 ms duration delivered at 1 Hz, administered using a BTX ECM 830 square pulse electroporator (Harvard Apparatus) via CUY650P3 3-mm round tweezer electrodes (NepaGene). DNA constructs were mixed in phosphate buffered saline (PBS) with Fast Green FCF dye (Sigma F7258-25G) to visualize injections. 0.75 μL of buprenorphine (1 mg/mL; ZooPharm LLC) was administered for post-operative pain control.

Constructs used for sparse labeling of L6CThNs for axon reconstruction were modified from the Supernova system^124^: *pTRE-DIO-FlpO* (15 ng/µL), a Cre-dependent Flp recombinase that expresses sparsely (https://www.addgene.org/118027/); and *pCAG-FSF-turboRFP-ires-tTA* (0.75 µg/µL), a Flp-dependent RFP. *pCAG-LSL-EGFP* (1.5 µg/µL), a Cre-dependent construct independent of the Supernova system^125^, was electroporated at higher concentrations to provide denser labeling of L6CThNs for layer identification and monitoring of electroporation efficiency. In experiments testing the effect of suppressed neuronal activity on L6CThN morphology, *pHSYN-FLEx-FRT-Kir2.1-2A-GFP* (1.5 ng/µL), a Flp-dependent expression construct for Kir2.1 with GFP labeling^126^ (https://www.addgene.org/161577/), was electroporated with *pTRE-DIO-FlpO* and *pCAG-FSF-turboRFP-ires-tTA*.

### Tamoxifen injections

*Lhx6-CreER;EGFP-Gria2* and *Lhx6-CreER;EYFP* pups received tamoxifen injections at P1, P2, and P3. Tamoxifen was dissolved in sunflower seed oil at a concentration of 2 mg/mL, and a uniform dose of 0.05 mL of solution (0.1 mg tamoxifen) was delivered intraperitoneally into the milk sac on each day.

### Stereotaxic injections

L6CThNs were labeled for electrophysiological recording using either Cre-dependent fluorescent reporter lines crossed to *Ntsr1-Cre* mice or with retrograde neuronal tracers. To retrogradely label L6CThNs, mice (P14-24) were anesthetized with ketamine (50 mg/kg, PennVet), dexmedetomidine (25 µg/kg, PennVet), and the inhalation anesthetic isoflurane (2.5-3%, PennVet) delivered in pure oxygen to an induction chamber. The mice were then administered atropine (0.065 mg/kg, PennVet). Anesthesia was maintained using 1-2.5% isoflurane in pure oxygen, and body temperature was maintained at 38-38.5°C. After shaving the head and disinfecting the surgical field, an incision was made to reveal the skull and a small craniotomy was made over the injection site using a dental drill. Approximately 90-135 nL of 1 mg/mL Alexa Fluor 555-conjugated cholera toxin subunit B (CTB-555; Invitrogen C34776) was pressure-injected into the ventral posterior medial nucleus of the thalamus (VPM; 1.1 mm posterior, 1.7 mm lateral, and 3.4 mm ventral relative to Bregma). Injections were slowly delivered via a glass pipette (inner diameter: 15-25 μm), which was left in place for 5 minutes. Skin incisions were sutured closed, and mice recovered under a heating lap with subcutaneous buprenorphine as a post-operative analgesic (0.05 mg/kg, PennVet). Injected mice were euthanized and used for slice electrophysiological recordings 1-7 days after stereotaxic injections of CTB.

### Electrophysiological recording

Mice aged P3-P24 were deeply anesthetized with isoflurane, decapitated, and brains were rapidly extracted in ice-cold sucrose-based cutting solution consisting of (in mM) 76 NaCl, 25 NaHCO_3_, 25 glucose, 75 sucrose, 2.5 KCl, 1.25 NaH_2_PO_4_, 0.5 CaCl_2_, 7 MgSO_4_ (pH 7.3, 315 mOsm). The hemisected forebrain was mounted on a 30° ramp and 300 µm parasagittal slices were cut in ice-cold sucrose solution using a vibratome (Leica VT-1200s). Brains from neonatal mice aged P3-5 were embedded in a gel of 4% low-melting-point agarose (Sigma A9045) in sucrose solution to stabilize tissue for cutting^132^. Agarose solution close to gelling temperature (∼35°C) was poured over the lateral side of the hemisected forebrain and cooled with cold sucrose before mounting on the cutting ramp. Slices were transferred to warm sucrose solution and incubated for 30 minutes at 32-35°C, then transferred to warm artificial cerebrospinal fluid (ACSF) and allowed to cool to room temperature. ACSF consisted of (in mM) 125 NaCl, 26 NaHCO_3_, 2.5 KCl, 1.25 NaH_2_PO_4_, 1 MgSO_4_, 20 D-(+)-glucose, 2 CaCl_2_, 0.4 ascorbic acid, 2 pyruvic acid, 4 L-lactic acid (pH 7.3, 315 mOsm). All solutions were continuously bubbled with a 95/5% O_2_/CO_2_ gas mixture. Brain slices were mounted on glass coverslips coated with poly-L-lysine (Sigma P4832) and visualized with an upright microscope (Zeiss AxioExaminer; 40x objective, 1.0 N.A) using infrared differential interference contrast (IR-DIC) and wide-field fluorescence microscopy. Slices containing the whisker-associated barrel region of primary somatosensory cortex and that had visible apical dendrites of infragranular pyramidal neurons approximately parallel to the plane of the slice were selected for recording.

Borosilicate glass pipettes (Sutter BF150-110-10) with a resistance of 2-4 MΩ were pulled using a horizontal pipette puller (Sutter P-97). For current-clamp recordings, pipettes were filled with a solution consisting of (in mM): 2.7 KCl, 120 KMeSO_3_, 9 HEPES, 0.18 EGTA, 4 MgATP, 0.3 NaGTP, 20 phosphocreatine(Na) (pH 7.3, 295 mOsm). For voltage-clamp recordings, pipettes were filled with a solution of (in mM): 115 CsMeSO_4_, 0.4 EGTA, 5.0 TEA-Cl, 1 QX314, 2.8 NaCl, 20 HEPES, 3.0 ATP magnesium salt, 0.5 GTP sodium salt, 10 phosphocreatine disodium salt, 0.1 spermine (pH 7.3, 295 mOsm). Voltage-clamp internal solutions contained spermine (0.1 mM, Millipore-Sigma S1141) for all recordings of GFP-positive neurons in *G42* mice and all recordings of EGFP or EYFP-positive neurons from *Lhx6-CreER;EGFP-Gria2* and *Lhx6-CreER;EYFP* mice to maintain rectification properties of AMPARs that are dependent on interaction with intracellular polyamines^100–102^. Spermine was included in all but 2 voltage-clamp recordings of L6CThNs labeled with *Ntsr1-Cre;tdTomato*. In most recordings, biocytin (0.25%, Sigma B4261) was included in the internal solution for *post hoc* visualization of recorded cells. Glass pipettes were mounted in pipette holders (Molecular Devices 1-HL-U) and lowered towards the targeted cell using a micromanipulator (Sutter MPC-200) while applying positive pressure to the pipette until reaching the surface of the cell. Negative pressure was then applied to generate a seal with a resistance greater than 1 GΩ. A brief burst of negative pressure was then used to break the membrane and attain whole-cell configuration. All recordings were performed in ACSF at ∼32-34° C.

Access resistance was measured from the magnitude of transient currents during a 5-mV voltage step. Only neurons with an access resistance less than 30 MΩ were included in subsequent analyses. An initial resting membrane potential was measured immediately upon transitioning to current clamp after breaking in, and intrinsic properties of neurons recorded in current clamp were assessed with 1 s depolarizing and hyperpolarizing current steps. Electrophysiological data was collected via a CV-7B headstage connected to a Multiclamp 700B amplifier (Molecular Devices) and an analog-digital converter (HEKA Instrutech ITC-18). Custom-written software in IGOR Pro (Wavemetrics) was used to record digitized data. All signals were low-pass filtered at 10 kHz and sampled at 20-100 kHz. Recordings were not compensated for liquid junction potential.

As the types of L6CThN and PV interneurons differ in the top and bottom of L6a^16,19,70^, for all paired recording experiments conducted within L6, only cells within the top 50% of L6a were targeted for recording, enriching the recordings for L6CThNs that project to VPM-only and for interlaminar PV interneurons. L6a boundaries were identified using the vertical extent of tdTomato-positive cell bodies in *Ntsr1-Cre;tdTomato* mice or retrogradely labeled L6CThNs following CTB-555 injections in the VPM of *Lhx6-CreER;EGFP-Gria2 and Lhx6CreER;EYFP* mice. For paired recordings in L4, the morphological appearance of the barrels under IR-DIC microscopy and the band of ChR2-EYFP fluorescence at the L4-L5a border in *Ntsr1-Cre;ChR2-EYFP* mice were used to identify layer location, and PV interneurons were selected using green fluorescence^20^. Neighboring excitatory neurons were selected based on cell body morphology under IR-DIC. In *Lhx6-CreER* mice with either the *EGFP-Gria2* or *EYFP* alleles, green fluorescence was used to target Lhx6 interneurons while red fluorescence from retrograde labeling with CTB-555 injection in the VPM was used to target L6CThNs.

### Analysis of intrinsic electrophysiological properties of parvalbumin-positive interneurons

Initial resting membrane potentials were calculated from the average of 15 traces acquired at 1 Hz shortly after obtaining the whole-cell current-clamp configuration. The input resistance (R_in_) was calculated using 1 s hyperpolarizing current steps of -25 to -100 pA. The properties of action potentials (APs) were calculated using the trace with the fewest action potentials evoked by a 1 s depolarization (L6: 7.7 ± 1.17 Hz mean ± SEM, n = 61 cells, N = 26 animals; L4: 9.1 ± 1.6 Hz mean ± SEM, n = 42 cells, N = 13 animals; Wilcoxon rank-sum test for difference in number of spikes used between layers, p = 0.705). AP threshold was calculated as the membrane potential at which dV_m_/dt exceeded 20 V/s. The AP half-width was calculated as the difference in time between when the membrane potential attained half-maximum amplitude in the rising versus the falling phases of the AP. Maximum rise rate of the AP was calculated as the maximum value of the first derivative of membrane potential during the rising phase of the AP. Maximum fall rate of the AP was calculated as the absolute value of the minimum value of the first derivative of the AP during the falling phase of the AP.

### Optogenetic stimulation of L6CThNs

L6CThN axons were photostimulated with blue light delivered through a 40x objective (∼470 nm wavelength, 200-250 µm diameter, 30-550 mW/mm^2^). To determine whether photostimulation efficiently elicits action potentials in slices from mice homozygous or hemizygous for *ChR2* across the age range we tested, whole-cell recordings of ChR2-EYFP-expressing L6CThNs were made, and ChR2 expression was confirmed with a 1 s pulse of blue light delivered over L6 while voltage clamping the neuron at -70 mV. Neurons that exhibited a steady-state inward current during 1 s photostimulation were then tested in current-clamp to determine whether photostimulation with 3 ms light pulses, delivered as a train of 10 pulses at 10 Hz, reliably induced action potentials. The number of action potentials elicited per 3 ms light pulse was quantified. To assess the responses of L4 PV and L4 excitatory neurons to activation of L6CThN synapses at different ages, mice homozygous for *ChR2-EYFP* were used. Pairs composed of a L4 PV and an unlabeled L4 neuron were assessed simultaneously using trains of five 3 ms pulses of light stimulation delivered at 10 Hz to control for potential differences across slices.

### Analysis of responses to photostimulation of L6CThN axons

Response amplitudes were quantified from the average of 30-100 trials. Pairs with fewer than 30 trials were not analyzed. For L4 PV interneurons, the averaged maximum excursion measured over 0.5 ms within 12.5 ms of photostimulation was determined. For unlabeled L4 neurons, the maximum excursion within 20 ms of photostimulation was determined as the latencies to response peaks were typically slower than for PV interneurons (Mean latency to peak response, L4 Excitatory neurons: P7-8: 6.5 ± 2.4 ms, P10-11: 10.7 ± 5.6 ms, P14-15: 13.2 ± 2.1 ms; L4 PV interneurons: P7-8: 6.1 ± 2.0 ms, P10-11: 7.1 ± 2.1 ms, P14-15: 5.4 ± 1.3 ms). Some cells at P7-8 and P10-11 exhibited slow depolarizing or hyperpolarizing responses with peaks outside of the 12.5 ms and 20 ms ranges (L4 Excitatory neurons: P7-8: 4/12, P10-11: 2/20; L4 PV interneurons: P7-8: 4/12, P10-11: 3/20). In these cases, the amplitude of the response was measured at 12-12.5 ms for PV interneurons and at 19.5 to 20 ms for unlabeled neurons. When no discernible peak occurred in response to photostimulation, the amplitude of the response was measured at the average quantification window used for identified peaks measured in other cells of the same type and age group.

### Testing for synaptic connectivity in paired recordings

When testing for unitary synaptic connections, pairs composed of two L6CThNs or a L6CThN and a PV interneuron in upper L6 were targeted for recording. The two cells were alternately stimulated every 10 s with a 20 Hz train of ten 0.5-1.5 nA current pulses, 3 ms in duration, to evoke action potentials while responses were recorded in current clamp in the other neuron. Only pairs in which both neurons fired regenerative APs with peak membrane potentials greater than 0 mV were analyzed. To prevent spontaneous spiking of more depolarized neurons, which was more common in younger animals, up to -50 pA of hyperpolarizing current was applied during connectivity tests.

To determine if L6CThNs form silent synapses onto either L6 PV interneurons or other L6CThNs during P6-P9, paired recordings were conducted with the candidate postsynaptic neuron recorded in voltage clamp, and the candidate presynaptic neuron recorded in current clamp as described above. Trains of APs were evoked in the candidate presynaptic cell as described above. In these pairs, testing for synaptic connectivity was performed in alternating blocks of 5, with the candidate postsynaptic neuron alternately held at -85 mV and +40 mV. Wait times of 10 s were imposed between tests at one membrane potential, and an additional 10 s wait time was added following each change in membrane holding potential between -85 mV and +40 mV.

### Analysis of paired recordings to determine connectivity

To determine if pairs of recorded neurons were monosynaptically connected, 50-150 postsynaptic responses were averaged. Traces where the membrane potential of either neuron exceeded -45 mV, depolarized more than 20 mV from the neuron’s initial membrane potential, or in which either neuron was spontaneously spiking, were not analyzed. If more than 5 traces in a row met these criteria, no further traces were analyzed from that pair. Pairs with fewer than 50 traces acquired were excluded from analysis. Pairs were determined to be connected if the average unitary EPSP or IPSP magnitude exceeded twice the value of the root-mean-square noise of the average membrane potential measured in a 200 ms period beginning 600 ms after the last presynaptic 3 ms current pulse. The uEPSP amplitude and the paired-pulse ratio (PPR), calculated as the magnitude of the second EPSP divided by the magnitude of the first EPSP, for each unitary synaptic connection were assessed using the first 25 responses collected.

In experiments in which the postsynaptic neuron was recorded in voltage clamp, 15-100 responses were averaged at each holding potential (-85 mV and +40 mV). Responses recorded at a holding potential of -85 mV with a leak current more negative than -300 pA or more than 100 pA less than the initial steady-state current at -85 mV, were excluded from analysis. Responses recorded at +40 mV with a leak current 100 pA more positive than the initial steady-state current at +40 mV, or that exhibited oscillatory active currents during presynaptic neuron stimulation were excluded from analysis. If more than 5 responses in a row at either holding potential did not satisfy these criteria for inclusion, no subsequent traces were included at either holding potential. Tested connections that did not have 15 qualifying responses recorded at both holding potentials were excluded. Tested potential connections were determined to be connected if the magnitude of the uEPSC exceeded five times the root-mean-square noise of the average membrane potential. The threshold used for determining connectivity in voltage clamp was higher than that used in current clamp due to the lower noise levels in voltage clamp relative to uEPSC amplitudes.

### Electrical stimulation of intracortical synaptic inputs

To test if GluA2 overexpression after tamoxifen injection in *Lhx6-CreER;EGFP-Gria2* mice reduced the proportion of calcium-permeable AMPA receptors at synapses onto Lhx6 interneurons, we assessed the current-voltage relationships of AMPA synaptic currents after electrical stimulation of the neuropil in L6. To isolate AMPA receptor-mediated synaptic responses, recordings were conducted with 5 µM RS-CPP (Tocris 0173) and 100 µM picrotoxin (Tocris 1128) in the external solution. Current-voltage relationships of synaptic inputs were assessed by clamping the neuron at holding potentials between -80 mV and +80 mV (20 mV intervals) for P14-P22 recordings, or between -60 mV and +60 mV (30 mV intervals) for P5-P7 recordings, in a pseudorandom order while stimulating with a bipolar electrode (FHC, MX21XES(DB9)). Stimulation intensity was controlled with an AMPI Iso-Flex stimulus isolator. Stimulation intensity for I-V curves was first set by gradually increasing stimulator voltage while holding the cell at -70 mV until an EPSC with a latency of approximately 2-5 ms was observed. Stimulus intensity ranged from 0.01 mA to 0.5 mA (duration: 0.1 ms). 5-10 repeats of the I-V curve were then acquired using a constant stimulation voltage.

### Analysis of responses to electrical stimulation

To determine the current-voltage relationship of synaptic currents elicited by intracortical electrical stimulation in neurons from *Lhx6-CreER;EYFP* and *Lhx6-CreER;EGFP-Gria2* mice, EPSC magnitude was calculated from the average of 5-10 trials at each holding potential. Neurons for which the leak current was more negative than -500 pA at a -70 mV holding potential were excluded from analysis. Only traces in which the hold current returned to the steady-state baseline within 1 s after stimulation were analyzed. Rectification index (RI) was calculated as the ratio of the absolute value of the average EPSC recorded at +80 mV to that recorded at -80 mV in P14-P22 animals. In P5-P7 animals, RI was calculated as the ratio of the average EPSC magnitude recorded at +60 mV to that at -60 mV. Liquid junction potential (∼11 mV) was not compensated for. Data were analyzed blind to genotype.

### Immunohistochemistry in fixed tissue sections

To visualize PV and SOM neurons and test for effects of GluA2-EGFP expression on the density and distribution of Lhx6 interneuron subtypes, we immunostained for PV and SOM in fixed slices of barrel cortex. For mice P7 or older, mice were deeply anesthetized with isoflurane and transcardially perfused with ice-cold phosphate buffered saline (PBS), followed by 4% paraformaldehyde (PFA) in PBS. Brains were extracted and post-fixed overnight in 4% PFA in PBS at 4°C. For neonates younger than P7, mice were deeply anesthetized with isoflurane or hypothermia, decapitated, and brains were directly dissected out and fixed overnight in 4% PFA in PBS at 4°C. Brains were washed 3x in PBS and 50 µm coronal sections of the barrel cortex were cut on a vibratome (Leica VT1000s). Sections were then washed 3x in PBS, blocked and permeabilized in a solution of 0.3% Triton X-100 and 3% Normal Donkey Serum (NDS) in PBS for 1 hour at RT, and incubated overnight at 4°C in blocking solution with primary antibodies at a concentration of 1:1000 each (rabbit anti-somatostatin, BMA T4103; mouse anti-parvalbumin, Sigma-Aldrich P3088; and chicken anti-GFP, Aves GFP-1020). Sections were then washed 3x in PBS and incubated for 90 minutes at RT in blocking solution with secondary antibodies at a concentration of 1:1000 each (donkey anti-rabbit Alexa Fluor 555, Invitrogen A31572; donkey anti-mouse 647, Invitrogen A31571; and donkey anti-chicken Alexa Fluor 488, Jackson 703-545-155). Finally, sections were washed 3x in PBS, counterstained with DAPI, and mounted with Aqua Polymount for imaging on a confocal microscope. For each section of the barrel cortex, a maximum intensity projection of 3 optical sections acquired on a Zeiss LSM800 confocal microscope with a 20X objective (NA 0.8) spanning a tissue depth of 5.7 µm was generated. From this maximum intensity projection, one or two regions forming a rectangle 400 µm-wide in the horizontal direction and spanning from the pia to the white matter in the vertical direction were selected for analysis. Experimenters, blinded to genotype, manually counted cells with DAPI-stained nuclei immunopositive for PV, SOM, EYFP or EGFP using FIJI/ImageJ with the Cell Counter plugin (https://imagej.net/ij/plugins/cell-counter.html).

### Tissue clearing and light-sheet microscopy for axonal reconstruction

Mice aged P4 or P7 were anesthetized on ice, and mice aged P10 or P14 were anesthetized using carbon dioxide. Animals were first transcardially perfused with a modified PHEM buffer that consisted of, in mM: 27 PIPES, 25 HEPES, 5 EGTA, 0.47 MgCl_2_, pH = 6.9, followed by ice-cold 4% PFA in PHEM with 5% sucrose, 0.1% Triton X-100 and 100 μg/mL heparin. Brains were dissected out and post-fixed overnight at 4°C in the same 4% PFA solution. Then, all samples were washed multiple times in PBS, and stored in PBS with 0.02% sodium azide until clearing.

Brain tissue was cleared for imaging using the solvent-based clearing protocol Adipo-Clear^133^. Briefly, brains were progressively dehydrated in methanol mixed with a buffer solution containing 0.3 M glycine and 0.1% Triton X-100 in water (pH 7.0), delipidated with 100% dichloromethane, and washed with 100% methanol. Samples were then rehydrated and stored in PTxwH buffer consisting of 0.1% Triton X-100, 0.05% Tween 20, and 2 µg/mL heparin in PBS. Immunohistochemistry was performed by incubating with rabbit anti-TagRFP (ThermoFisher R10367) and chicken anti-GFP (Aves GFP-1020) antibodies at 1:1000 in PTxwH buffer for 4-5 days at 37°C. Brains were washed multiple times in PTxwH buffer and secondary antibody labeling was performed under the same conditions as primary antibody labeling, with goat anti-chicken Alexa Fluor 488 and goat anti-rabbit Alexa Fluor 555 antibodies (Jackson ImmunoResearch) at 1:1000. TO-PRO-3 iodide was added to the secondary antibody solution to label nuclei. After immunohistochemistry, brains were dehydrated using a progressive methanol/water gradient, incubated with dichloromethane for 1 hour, and incubated with dibenzyl ether for 2 days to ensure refractive index matching. After incubation with dibenzyl ether, samples were transferred to ethyl cinnamate for long-term storage and imaging. Samples were imaged using a Zeiss Lightsheet 7 light-sheet microscope with a 20X (NA 1.0, nd = 1.53) objective at 0.8x zoom.

### Neuronal morphological reconstruction in cleared brains

Reconstruction of the axonal morphology of L6CThNs was performed in FIJI/ImageJ^130^ using the SNT plugin^131^ (https://imagej.net/plugins/snt/). Axon and dendrite tracing was performed in semi-automatic configuration using the A* search algorithm on the tRFP staining channel. Only neurons labeled with both tRFP and either the control EGFP or Kir2.1-EGFP were reconstructed. Each reconstruction was verified by two expert tracers. Reconstructions were exported from SNT in .swc format, and custom-written software in MATLAB was used to quantify axon morphology. Neurite length and branch point location was quantified with a resolution of 0.285 x 0.285 x 0.655 µm, the minimum voxel size. Branch points were located at each bifurcation of the axon or dendrite. Each continuous segment of axon or dendrite between two branch points or between a branch point and an end point was considered an axon or dendrite branch for quantification purposes. All branches shorter than 10 µm were removed from the analysis to ensure that branch quantifications did not include small axonal protrusions, and to minimize the effects of imaging artifacts. Only the intracortical portions of the axon reconstructions above the white matter border were quantified.

Cortical boundaries at the pial surface and white matter (WM) as well as boundaries between cortical layers were determined using images of the TO-PRO-3 signal or EGFP signal in *Ntsr1-Cre* mice electroporated with a Cre-dependent *EGFP* or *Kir2.1-EGFP* construct. The intersections of the radial axis of the cell with the pia and WM were manually annotated, and axon and dendrite reconstructions were rotated and aligned to this pia-WM axis for spatial analyses. To segment the cortical volume into layers, cortical layer boundaries were manually annotated using points 50-100 μm apart in the x-y plane, on every 100th z-section (65 μm apart in z) and 3-dimensional surfaces were fit to each set of points using a paraboloid function (y= a*x*^2^ + b*z*^2^ + c*xz* + d*x* + e*z* + f) with custom-written software in Python 3.9.0. Axon and dendrite reconstructions were segmented using their location relative to layer boundary surfaces, and length and branch point location were quantified within each cortical layer. Views of axons or dendrites in the tangential plane were generated by projecting the layer-segmented neurites of multiple cells onto a plane that was perpendicular to the radial axis of the cell and roughly parallel to the pial and WM surfaces. The number of cells shown in each tangential view was determined by the smallest experimental group in each comparison, and the cells closest to the mean axon or dendrite length within the layer of interest for each experimental group are shown. Horizontal area spanned within each layer was calculated by binning points in the axon or dendrite reconstruction within each layer, then defining a 2-dimensional convex hull in the horizontal plane by identifying a minimal set of points that, when connected by line segments, enclosed the arbor’s entire horizontal extent in each layer. The points comprising the convex hull and the area of the shape it defined were calculated using the convhull function in MATLAB.

### Quantification and statistical analysis

Data visualization, analysis, and statistical comparisons were performed in MATLAB (Mathworks). Image acquisition and stitching were performed in Zen Blue (Zeiss), and image processing was performed in FIJI/ImageJ. Statistical details such as the number of samples, number of mice, and statistical tests used are reported in the figure legends. Non-parametric tests for differences in median such as Wilcoxon rank-sum tests and Wilcoxon signed-rank tests were used to compare continuous data between groups, and parametric tests such as ANOVA were only used after standardized data residuals were compared to a standard normal distribution using a Kolmogorov-Smirnov test and found to be not significantly different. Chi-square tests of independence and Fisher’s exact tests were used to compare nominal data between groups. Permutation tests with 10,000 repeats were used to determine if correlations between continuous variables were statistically significant. Where applicable, the Bonferroni correction for multiple comparisons was applied to the standard value for statistical significance of alpha = 0.05. All error bars represent the standard error of the mean unless otherwise specified.

