## Supplementary material for "Cell-type-selective synaptogenesis during the development of excitatory connectivity in the mammalian neocortex": GutmanWei, Sudarsanam et al Supplemental Figures and Tables

### SUPPLEMENTAL INFORMATION:

**Figures S1-S7 and Tables S1-S2**

**Cell-type-selective synaptogenesis during the development of excitatory connectivity in the mammalian neocortex**


Alan Y. Gutman-Wei*, Sriram Sudarsanam*, Alec G. Cabalinan, Naseer Shahid, Anny Shi, Luis E. Guzman Clavel, Sophia M. Spindler-Krage, Amit Agarwal, Alex L. Kolodkin, Solange P. Brown[1]

[1]Lead contact.
*Co-first authors.

Correspondence should be addressed to:

Solange P. Brown


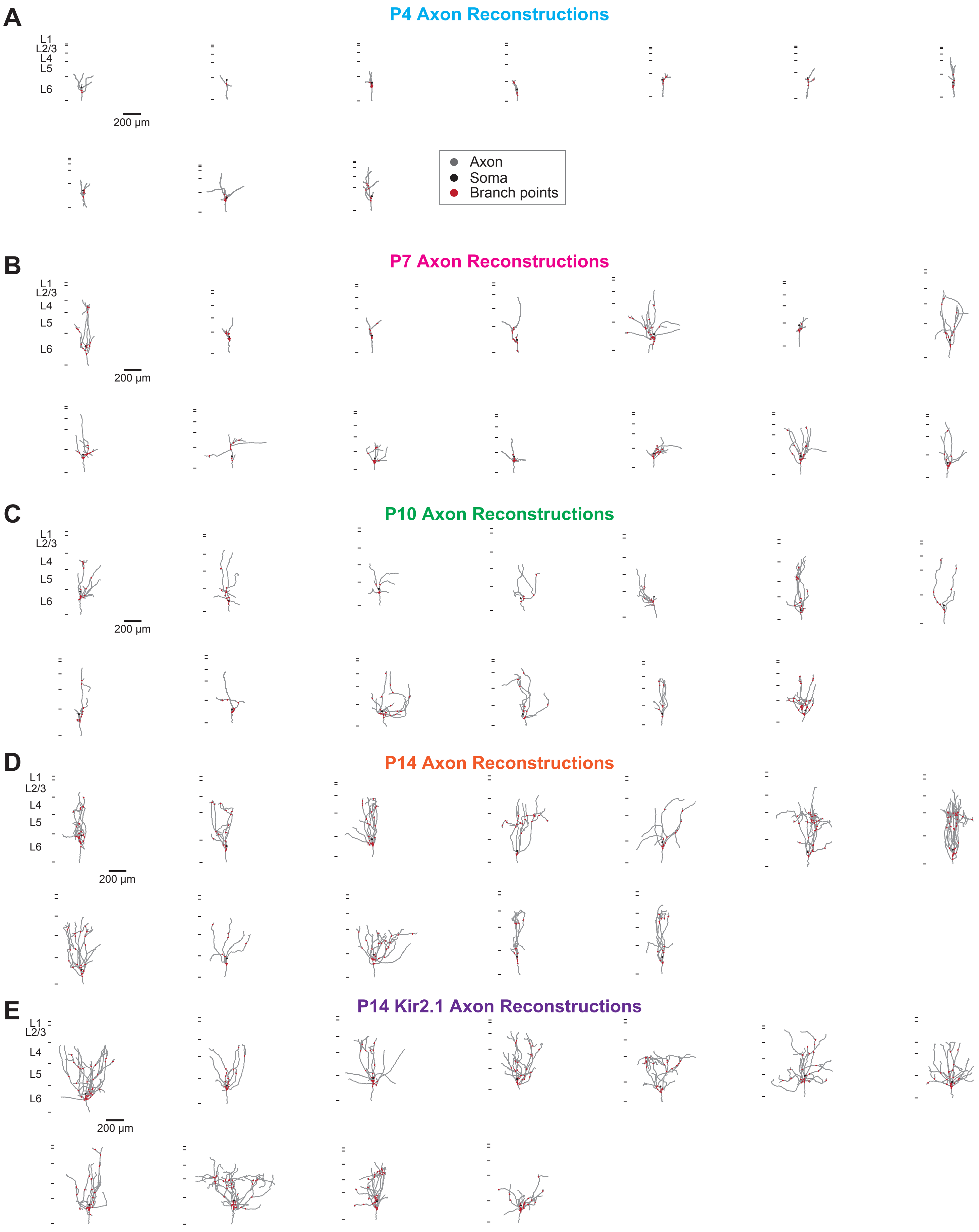


Figure S1, Gutman-Wei et al. 2025

**Figure S1. Reconstructions of the intracortical axons of layer 6 corticothalamic neurons (L6CThNs) across the first two postnatal weeks. Related to Figures 1, 2 and 3. (A-D)** Axonal reconstructions of all L6CThNs traced from postnatal day 4 (P4, A, n = 10 cells, N = 3 animals), P7 (B, n = 14 cells, N = 5 animals), P10 (C, n = 12 cells, N = 5 animals) and P14 (D, n = 12 cells, N = 8 animals) mice. Red points represent axon branch points; black circles represent soma locations. **(E)** Axonal reconstructions of all Kir2.1-expressing L6CThNs traced from P14 *Ntsr1-Cre* mice electroporated with Cre-dependent *Kir2.1* constructs (n = 11 cells, N = 7 animals). Scale bars: 200 μm.

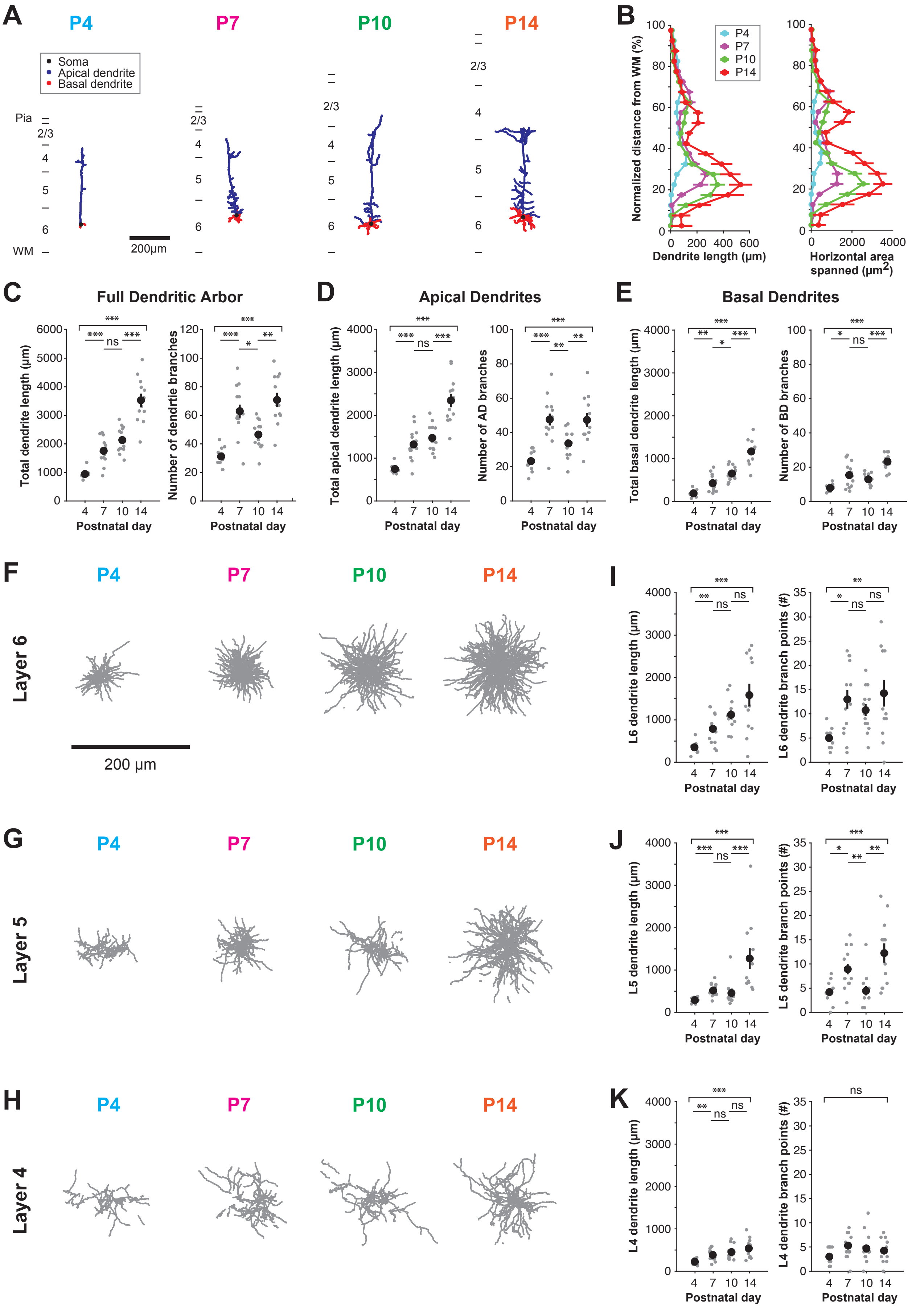


Figure S2, Gutman-Wei et al. 2025

**Figure S2: Dendritic development of layer 6 corticothalamic neurons (L6CThNs). Related to Figures 1 and 2. (A)** Example dendritic reconstructions of L6CThNs from postnatal day 4 (P4), P7, P10, and P14 mice. Example cells correspond to those reconstructed in the top row of Figure 1B. Blue represents the apical dendrites, red the basal dendrites, and black the soma. **(B)** Average dendritic length distribution across the depth of the cortex, measured at 5% intervals (left), and dendritic area spanned in the horizontal plane (right) at each cortical depth at P4, P7, P10 and P14 (P4: n = 10 cells, N = 3 animals; P7: n = 14 cells, N = 5 animals; P10: n = 12 cells, N = 5 animals; and P14: n = 12 cells, N = 8 animals). **(C)** Total dendritic length (left) and total number of branches (right) at P4, P7, P10, and P14. Black circles represent the mean; grey circles represent individual cells (length $p < 10^{-7}$, branch number, $p < 10^{-5}$; Kruskal-Wallis tests). **(D)** Total length of L6CThN apical dendrites (left) and total number of apical dendrite branches (right) at P4, P7, P10, and P14 (length: $p < 10^{-6}$, branch number: $p < 10^{-4}$; Kruskal-Wallis tests). **(E)** As in (D), but for L6CThN basal dendrites (length: $p < 10^{-7}$; branch number: $p < 10^{-5}$; Kruskal-Wallis tests). **(F-H)** Views in tangential plane of the dendrites of L6CThNs in layer 6 (L6; F), layer 5 (L5; G), and layer 4 (L4; H). For each postnatal age and layer, the 10 cells with dendritic lengths within that layer closest to the mean of that layer and age are shown. **(I-K)** Total dendritic length and number of branch points of L6CThN dendrites in L6 (I), L5 (J) and L4 (K) across four age groups ( (I) $p < 0.009$ for all metrics; (J) $p < 0.0003$ for all metrics; (K) length: $p < 0.0003$, branch points: $p = 0.195$; Kruskal-Wallis tests). C,D,E,I,J,K: Pairwise Wilcoxon rank-sum tests with a Bonferroni correction for multiple comparisons were used to compare between consecutive age groups. *, $p < 0.05$; **, $p < 0.01$; ***, $p < 0.001$. Data in B-E and I-K shown as mean ± SEM. Scale bars: A, F-H: 200 μm.

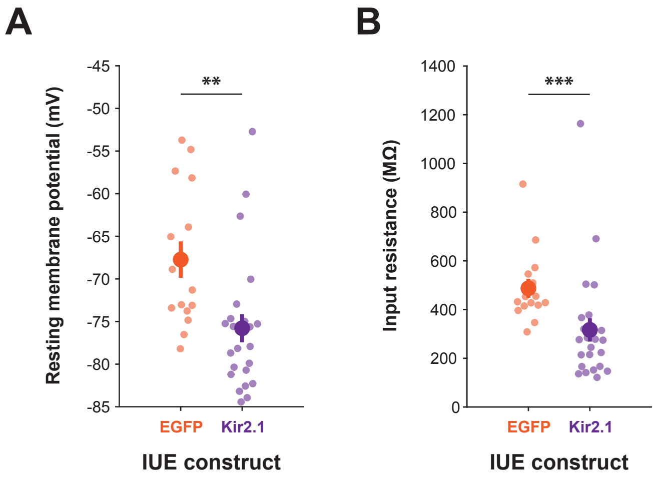


Figure S3, Gutman-Wei et al. 2025

**Figure S3: Expression of Kir2.1 decreases the excitability of layer 6 corticothalamic neurons (L6CThNs). Related to Figure 3. (A)** Resting membrane potentials of control L6CThNs expressing EGFP (n = 15 cells, N = 3 animals) and L6CThNs expressing Kir2.1 (n = 23 cells, N = 7 animals) in slices from postnatal day 6-7 (P6-7) mice ($p < 0.002$, Wilcoxon rank-sum test). **(B)** Input resistances of control L6CThNs expressing EGFP and L6CThNs expressing Kir2.1 in slices from P6-7 mice (, $p < 10^{-3}$, Wilcoxon rank-sum test). Data shown as mean ± SEM.

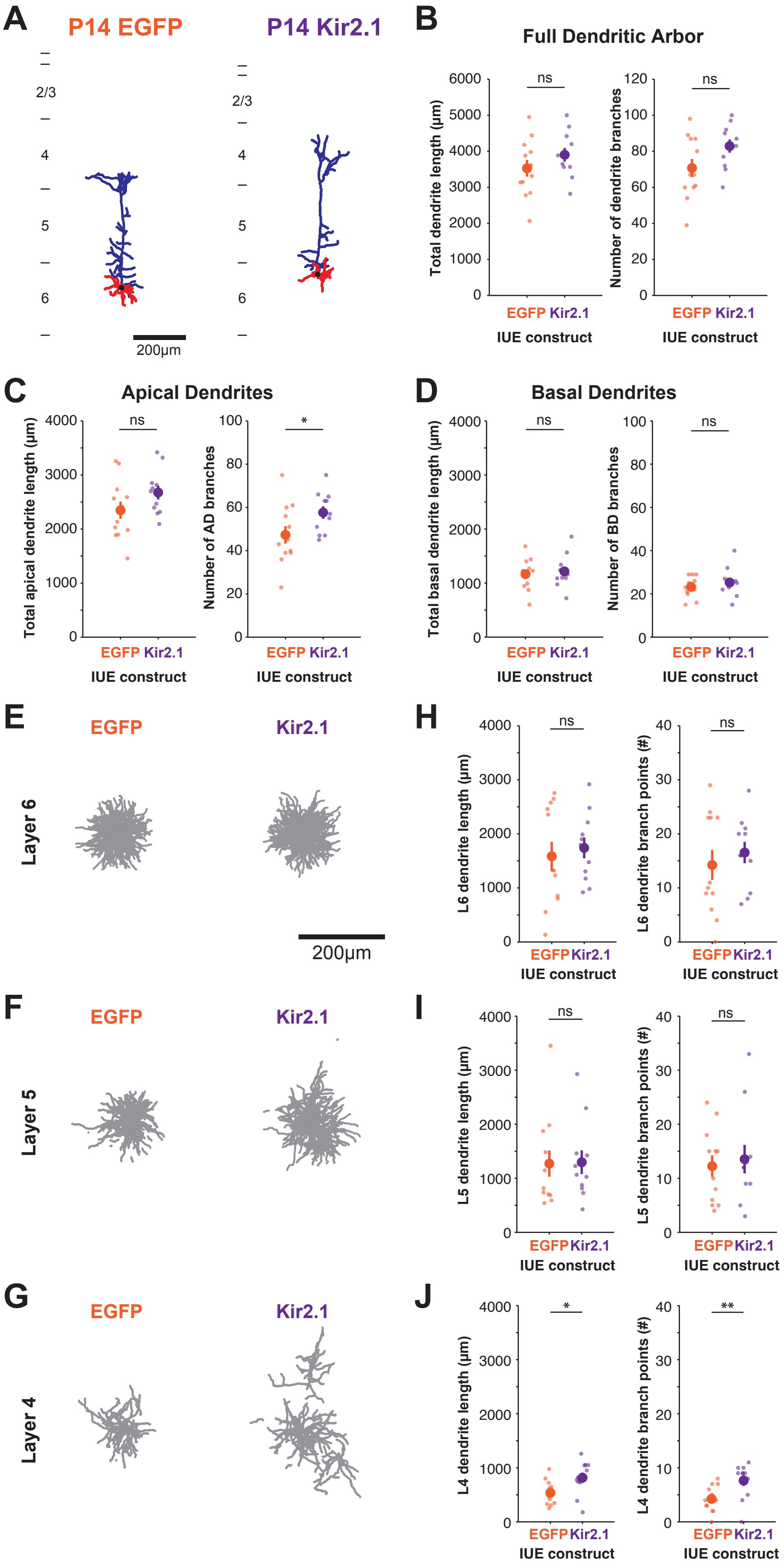


Figure S4, Gutman-Wei et al. 2025

**Figure S4: Kir2.1 overexpression results in increased branching of L6CThN apical dendrites in L4 at P14. Related to Figure 3. (A)** Example dendritic reconstructions of L6CThNs expressing control EGFP (left) and Kir2.1 (right). Blue represents the apical dendrite, red the basal dendrites, and black the soma. **(B)** Total dendritic length (left) and number of dendritic branches (right) for L6CThNs expressing control EGFP or Kir2.1. Large symbols represent the mean; small symbols represent individual cells (Length: $p = 0.281$, Branch number: $p = 0.09$, Wilcoxon rank-sum tests; Control EGFP: $n = 12$ cells, $N = 8$ animals; Kir2.1: $n = 11$ cells, $N = 7$ animals;). **(C)** Total length of the apical dendrites (left) and total number of apical branches (right) for L6CThNs expressing EGFP control or Kir2.1. Large symbols represent the mean; small symbols represent individual cells (Length: $p = 0.103$; Branch number: $p < 0.036$, Wilcoxon rank-sum tests). **(D)** As in (C), but for basal dendrites (Length: $p = 0.878$; Branch number: $p = 0.578$, Wilcoxon rank-sum tests). **(E-G)** Views in tangential plane of the dendrites of L6CThNs in layer 6 (L6; E), layer 5 (L5; F), and layer 4 (L4; G). For each experimental group and layer, the 11 L6CThNs with dendritic lengths in that layer closest to the mean of that layer and experimental group are shown. **(H-J)** Total dendritic length and total number of branch points for L6CThN dendrites in L6 (H), L5 (I), and L4 (J) from neurons expressing EGFP control or Kir2.1 ( (H) Length: $p < 0.689$, Branch points: $p < 0.829$; (I) Length: $p = 0.735$, Branch points: $p = 1$; (J) Length: $p < 0.021$, Branch points: $p < 0.008$, Wilcoxon rank-sum tests). Data in B-D, H-J shown as mean ± SEM.

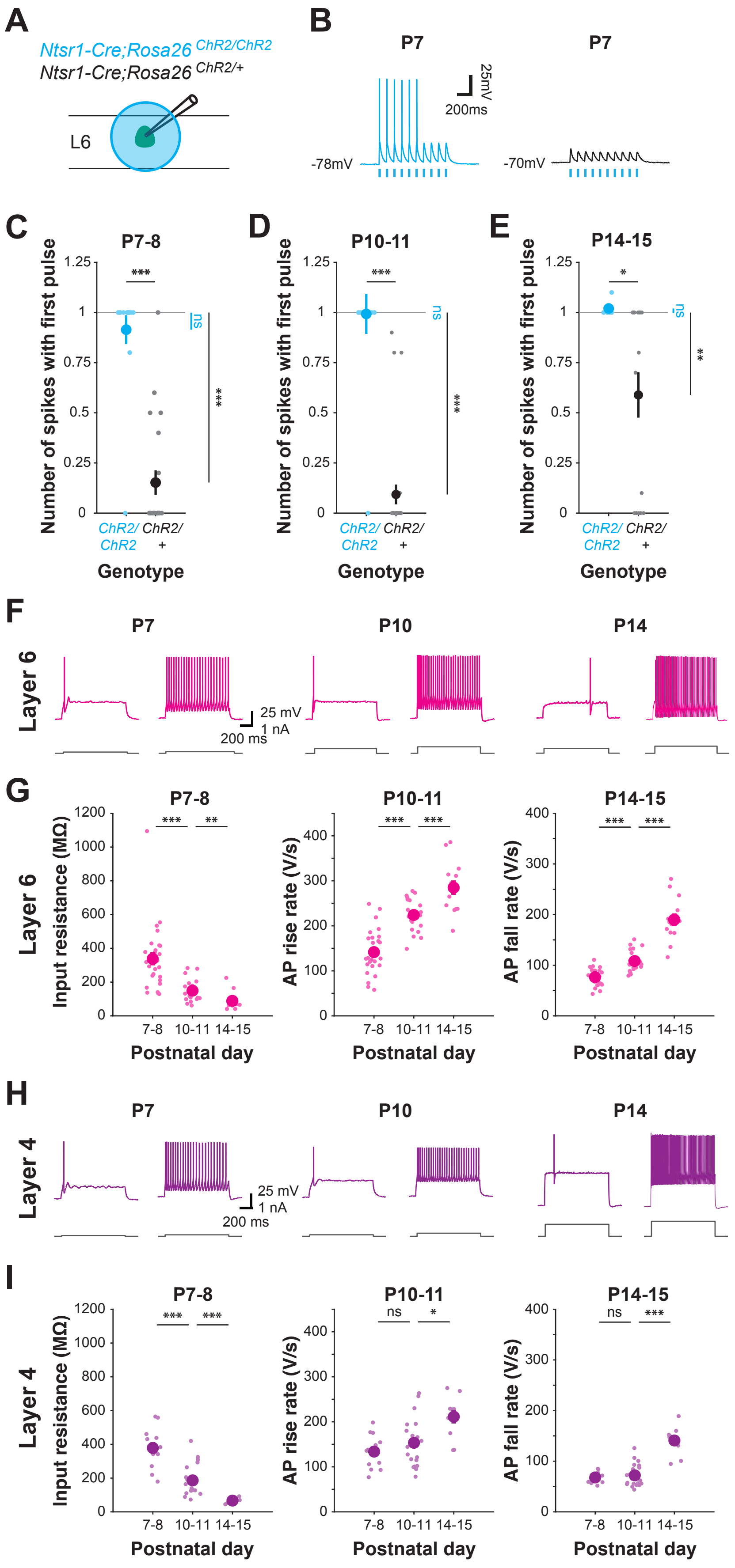


Figure S5, Gutman-Wei et al. 2025

**Figure S5. The effects of maturation on optogenetic activation of L6CThNs in neonates hemizygous or homozygous for *channelrhodopsin-2* (*ChR2*) and development of action potential properties in parvalbumin (PV) inhibitory interneurons in layer 6 (L6) and layer 4 (L4). Related to Figure 5. (A)** Experimental configuration for optogenetic photostimulation of L6CThNs in L6. **(B)** Example responses of a L6CThN from a mouse hemizygous for *Ntsr1-Cre* and homozygous (left) or hemizygous (right) for *ChR2* to optogenetic stimulation (3 ms light pulses at 10 Hz, cyan bars). **(C-E)** Average number action potentials elicited by the first light pulse in L6CThNs from mice homozygous (blue) or hemizygous (gray) for *ChR2* at P7-P8 (C), P10-11 (D), and P14-15 (E). Large symbols represent group means and small symbols represent the average of 10 responses of individual cells (C: n = 14 cells, N = 3 homozygous mice; n = 21 cell, N = 5 hemizygous mice; D: n = 14 cells, N = 3 homozygous mice; n = 28 cells, N = 7 hemizygous mice; E: n = 5 cells, N = 1 homozygous mouse; n = 18 cells, N = 3 hemizygous mice). C-E: Wilcoxon rank-sum tests were used to test for differences between genotypes, and Wilcoxon signed-rank tests were used to test for differences from 1 spike per pulse (*, $p < 0.05$; **, $p < 0.01$; ***, $p < 0.001$). **(F)** Example responses of a L6 PV interneuron (top) to 1 s depolarizing current pulses (bottom) at postnatal day (P) 7 (left), P10 (middle), and P14 (right). Responses to current pulses that were near rheobase (left) and that activated continuous spiking for a 1 s period (right) are shown. The P10 example response shown is from the same neuron displayed in Figure 5F. **(G)** Input resistance (left), the maximum rate of action potential rise (middle, AP rise) and the maximum absolute value of the rate of action potential fall (right, AP fall) for L6 PV interneurons measured at P7-P8, P10-P11, and P14-P15. Large symbols represent group means; small symbols represent individual cells (Input resistance: $p < 10^{-7}$; AP rise: $p < 10^{-8}$; AP fall: $p < 10^{-9}$, Kruskal-Wallis test for change between ages; P7-P8: n = 26 cells, N = 12 mice; P10-P11: n = 22 cells, N = 9 mice; P14-P15: n = 13 cells, N = 5 mice). **(H)** As in F, but example responses of L4 PV interneurons to 1 s depolarizing current pulses at P7, P10, and P14. The P10 example response shown is from the same neuron displayed in Figure

5G. **(I)** Input resistance, AP rise and AP fall for L4 PV interneurons measured at P7-P8, P10-P11, and P14-P15. Large symbols represent group means; small symbols represent individual cells (Input resistance: $p < 10^{-6}$; AP rise rate: $p < 0.007$; AP fall rate: $p < 10^{-4}$, Kruskal-Wallis test for change between ages; P7-P8: n = 12 cells, N = 4 mice; P10-P11: n = 20 cells, N = 5 mice; P14-P15: n = 10 cells, N = 4 mice). G,I: Pairwise Wilcoxon rank-sum tests with a Bonferroni correction for multiple comparisons were used to compare between consecutive age groups. *, $p < 0.05$; **, $p < 0.01$; ***, $p < 0.001$. Data in C-E, G and I shown as mean ± SEM.

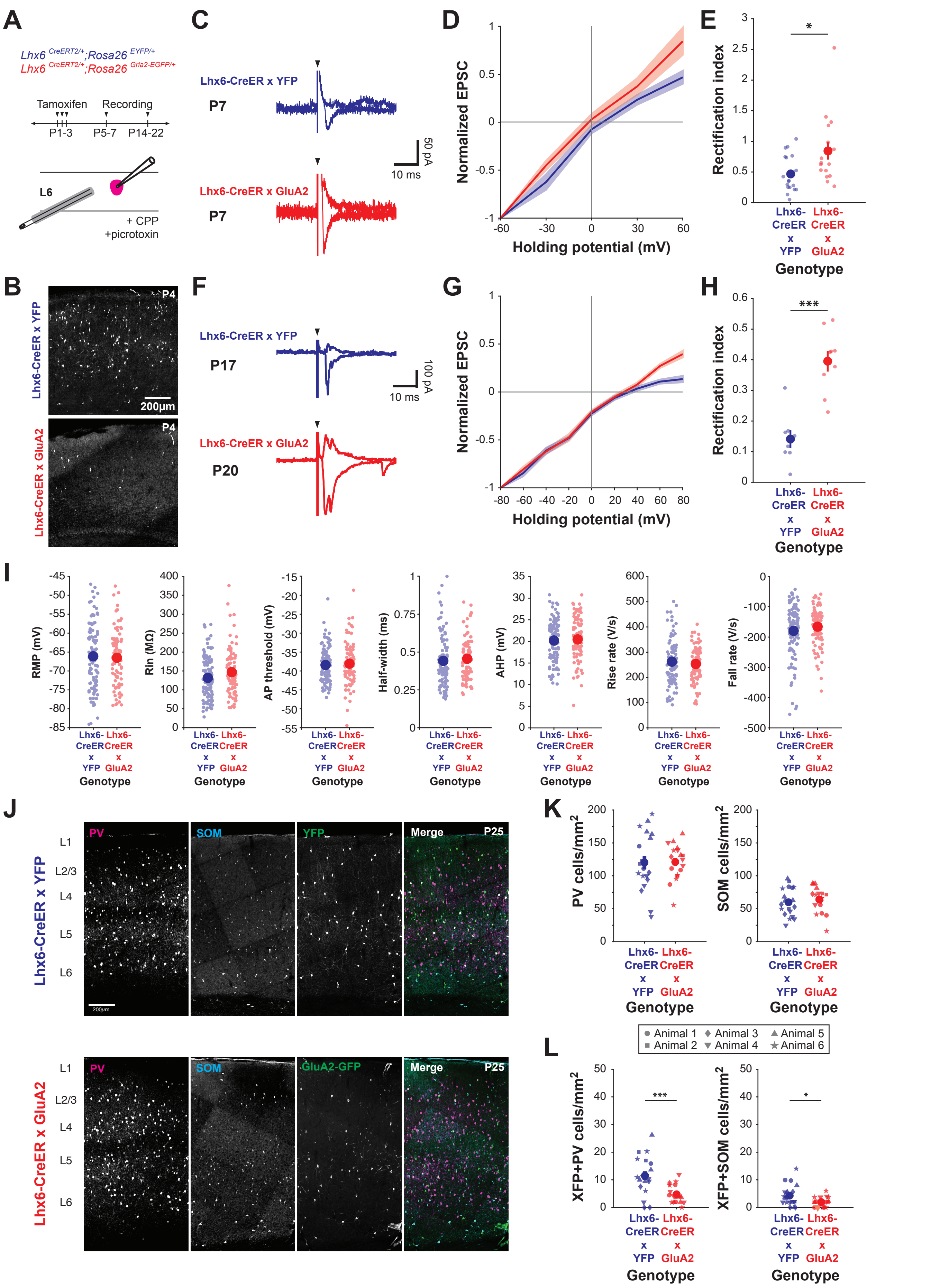


Figure S6, Gutman-Wei et al., 2025

**Figure S6. Cre-dependent expression of GluA2-EGFP in *Lhx6-CreER* mice alters synaptic current-voltage relationships but not the intrinsic electrophysiological properties or cell distribution of MGE-derived inhibitory interneurons. Related to Figure 6. (A**) Experimental paradigm, showing neonatal tamoxifen injections to induce transgene expression and the recording configuration for electrical stimulation of layer 6 corticothalamic neurons (L6CThNs). **(B)** Confocal images of barrel cortex in a coronal section processed for the immunohistochemical detection of fluorescent protein from a control postnatal day 4 (P4) *Lhx6-CreER;EYFP* mouse (top) and *Lhx6-CreER;EGFP-Gria2* mouse (bottom) following tamoxifen injections at P1, P2, and P3 (Green: EYFP/EGFP; Blue: DAPI counterstain). **(C)** Example average excitatory postsynaptic currents (EPSCs) recorded in fluorescent interneurons in slices from P5-P7 *Lhx6-CreER;EYFP* (top, blue) or *Lhx6-CreER;EGFP-Gria2* (bottom, red) mice following electrical stimulation of the adjacent neuropil in the presence of CPP and picrotoxin. Traces acquired at +60 mV and -60 mV are shown for each cell; the truncated stimulation artifact is indicated by a black arrowhead. **(D)** Current-voltage (I-V) curves for EPSCs recorded in fluorescent inhibitory interneurons in slices from P5-P7 *Lhx6-CreER;EYFP* (blue) and *Lhx6-CreER;EGFP-Gria2* (red) mice sampled at 30 mV intervals from membrane potentials of -60 mV to +60 mV. Thick lines show the average I-V curve for each genotype; shaded area shows the standard error of the mean for each genotype (*Lhx6-CreER;EYFP*: n = 19 cells, N = 7 mice; *Lhx6-CreER;EGFP-Gria2*: n = 16 cells, N = 9 mice). **(E)** Rectification indices of fluorescent interneurons in slices from P5-P7 *Lhx6-CreER;EYFP* (blue) and *Lhx6-CreER;EGFP-Gria2* (red) mice. Large symbols represent means; small symbols represent individual cells ($p < 0.0214$; Wilcoxon rank-sum test). **(F)** As in C, but for example responses recorded at +80 mV and -80 mV from fluorescent interneurons in slices from P16-P22 mice. **(G)** As in D, but for EPSCs recorded in fluorescent inhibitory interneurons in slices from P16-P22 mice. EPSCs were sampled at 20 mV intervals from membrane potentials of -80 mV to +80 mV (*Lhx6-CreER;EYFP*: n = 8 cells, N = 3 mice; *Lhx6-CreER;EGFP-Gria2*: n = 9 cells, N = 4 mice). **(H)** As in E, but for P14-P22 mice ($p < 0.0003$, Wilcoxon rank-sum test). **(I)** Intrinsic electrophysiological properties of fluorescent interneurons in layer 6 of *Lhx6-CreER;EYFP* (blue) and *Lhx6-Cre;EGFP-Gria2* (red) mice. Large symbols represent means; small symbols represent individual cells (Wilcoxon rank-sum tests for difference between genotypes: Resting membrane potential (RMP), p = 0.746; Input resistance ($R_{in}$), p = 0.097; Action potential (AP) threshold, p = 0.585; AP half-width, p = 0.263; After-hyperpolarization (AHP), p = 0.676; AP rise rate, p = 0.787; AP fall rate, p = 0.454; n = 106 cells, N = 35 *Lhx6-CreER;EYFP* mice; n = 91 cells, N = 39 *Lhx6-CreER;EGFP-Gria2* mice). **(J)** Example confocal images of parvalbumin (PV, far left), somatostatin (SOM, left) and green fluorescent interneurons (right) detected by immunohistochemistry in sections of the somatosensory cortex from *Lhx6-CreER;EYFP* (top row) and *Lhx6-CreER;EGFP-Gria2* (bottom row) mice. The three images are overlaid for comparison (far right). **(K)** Summary data comparing the densities of PV (left) and SOM (right) neurons in the barrel cortex of *Lhx6-CreER;EYFP* (blue) and *Lhx6-CreER;EGFP-Gria2* (red) mice aged P22-25. Large symbols represent the mean. Small symbols represent single regions of interest (ROIs). Symbol shapes indicate ROIs from the same individual animal within each genotype (n = 19 ROIs, N = 6 *Lhx6-CreER;EYFP* mice; n = 19 ROIs, N = 6 *Lhx6-CreER;EGFP-Gria2* mice; two-way ANOVA with mouse nested within genotype and litter, main effect of genotype: p = 0.7576 for PV density and p = 0.5315 for SOM density)**. (L)** As in K, but for the densities of fluorescent PV (left) and SOM (right) neurons from *Lhx6-CreER;EYFP* (blue) and *Lhx6-CreER;EGFP-Gria2* (red) mice (two-way ANOVA, $p < 0.001$ for PV and $p < 0.025$ for SOM interneurons). Data in E, H, I, K and L shown as mean ± SEM. Scale bars: B,J: 200 µm.

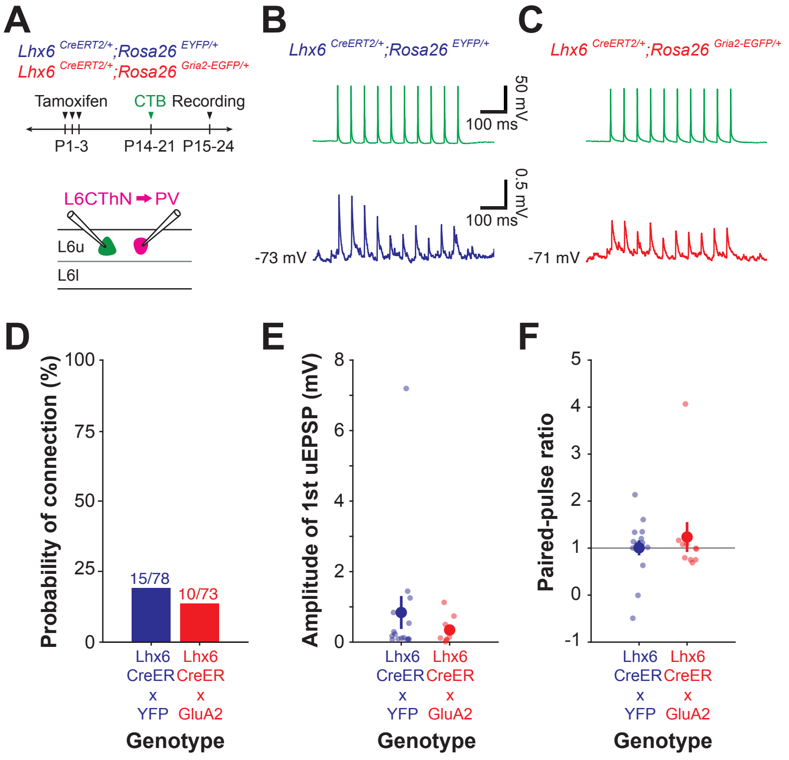


Figure S7, Gutman-Wei et al. 2025

**Figure S7. Reducing calcium-permeable AMPA receptors via overexpression of GluA2 in Lhx6 interneurons does not affect L6CThN→Lhx6 connectivity in L6. Related to Figure 6.** **(A)** Experimental paradigm, showing neonatal tamoxifen injections to induce transgene expression, labeling of layer 6 corticothalamic neurons (L6CThNs) with injections of the retrograde tracer, CTB, in the ventral posteromedial nucleus (VPM) of the thalamus, and the recording configuration. **(B)** Average unitary excitatory postsynaptic potential (uEPSP) recorded in a Lhx6 interneuron (bottom trace) in response to action potentials initiated in a presynaptic L6CThN (top, single trace, 20 Hz) from an *Lhx6-CreER;EYFP* mouse. **(C)** As in B for a L6CThN→Lhx6 synapse from an *Lhx6-CreER;EGFP-Gria2* mouse. **(D)** Probability of connection for tested L6CThN→Lhx6 connections from *Lhx6-CreER;EYFP* and *Lhx6-CreER;EGFP-Gria2* mice ($p = 0.39$, Fisher's exact test; *Lhx6-CreER;EYFP*: n = 78 pairs; N = 33 mice; *Lhx6-CreER;EGFP-Gria2*: n = 73 pairs; N = 38 mice). **(E)** Summary data comparing the average amplitudes of the first uEPSP for detected L6CThN→Lhx6 connections from *Lhx6-CreER;EYFP* and *Lhx6-CreER;EGFP-Gria2* mice ($p = 0.677$, Wilcoxon rank-sum test; *Lhx6-CreER;EYFP*: n = 15 connections, N = 14 animals; *Lhx6-CreER;EGFP-Gria2*: n = 10 connections, N = 9 animals). **(F)** Summary data comparing the paired-pulse ratio for detected unitary L6CThN→Lhx6 connections from *Lhx6-CreER;EYFP* and *Lhx6-CreER;EGFP-Gria2* mice ($p = 0.39$, Wilcoxon rank-sum test). Data in E and F shown as mean ± SEM.

### Table S1

| Postnatal Age (days) | L6CThN→PV Tested Connections | L6CThN→PV Detected Connections | L6CThN→L6CThN Tested Connections | L6CThN→L6CThN Detected Connections |
|---|---|---|---|---|
| **P3-4** | 14 | 0 (0%) | 28 | 0 (0%) |
| **P5-6** | 12 | 2 (16.7%) | 46 | 1 (2.2%) |
| **P7-8** | 17 | 1 (5.9%) | 42 | 2 (4.8%) |
| **P9-10** | 17 | 4 (23.5%) | 44 | 0 (0%) |
| **P11-12** | 14 | 6 (42.9%) | 24 | 0 (0%) |
| **P13-15** | 20 | 8 (40%) | 72 | 2 (2.8%) |

**Table S1:** Summary of unitary synaptic connections tested and detected for connections from layer 6 corticothalamic neurons (L6CThNs) to layer 6 parvalbumin-positive (L6 PV) interneurons and from L6CThNs to L6CThNs.

### Table S2

| Postnatal Age (days) | PV→L6CThN Tested Connections | PV→L6CThN Detected Connections |
|---|---|---|
| **P3-4** | 14 | 0 (0%) |
| **P5-6** | 12 | 0 (0%) |
| **P7-8** | 17 | 2 (11.8%) |
| **P9-10** | 17 | 8 (47.1%) |
| **P11-12** | 14 | 10 (71.4%) |
| **P13-15** | 20 | 13 (65%) |

**Table S2:** Summary of unitary synaptic connections tested and detected for connections from layer 6 parvalbumin-positive (L6 PV) interneurons to layer 6 corticothalamic neurons (L6CThNs).
